# Cohesin-mediated chromatin remodeling controls the differentiation and function of conventional dendritic cells

**DOI:** 10.1101/2024.09.18.613709

**Authors:** Nicholas M. Adams, Aleksandra Galitsyna, Ioanna Tiniakou, Eduardo Esteva, Colleen M. Lau, Jojo Reyes, Nezar Abdennur, Alexey Shkolikov, George S. Yap, Alireza Khodadadi-Jamayran, Leonid A. Mirny, Boris Reizis

**Affiliations:** Department of Pathology, New York University Grossman School of Medicine, New York, NY 10016, USA; Institute for Medical Engineering and Science, Massachusetts Institute of Technology, Cambridge, MA 02139, USA; Applied Bioinformatics Laboratories, New York University Grossman School of Medicine, New York, NY 10016, USA; Department of Microbiology and Immunology, Cornell University College of Veterinary Medicine, Ithaca, NY 14853, USA; Center for Immunity and Inflammation, Department of Medicine, New Jersey Medical School, Rutgers University, Newark NJ 07101, USA; Department of Genomics and Computational Biology, University of Massachusetts Chan Medical School, Worcester, MA 01605, USA; Department of Systems Biology, University of Massachusetts Chan Medical School, Worcester, MA 01605, USA; independent researcher; Department of Physics, Massachusetts Institute of Technology, Cambridge, MA 02139, USA

## Abstract

The cohesin protein complex extrudes chromatin loops, stopping at CTCF-bound sites, to organize chromosomes into topologically associated domains, yet the biological implications of this process are poorly understood. We show that cohesin is required for the post-mitotic differentiation and function of antigen-presenting dendritic cells (DCs), particularly for antigen cross-presentation and IL-12 secretion by type 1 conventional DCs (cDC1s) *in vivo*. The chromatin organization of DCs was shaped by cohesin and the DC-specifying transcription factor IRF8, which controlled chromatin looping and chromosome compartmentalization, respectively. Notably, optimal expression of IRF8 itself required CTCF/cohesin-binding sites demarcating the *Irf8* gene. During DC activation, cohesin was required for the induction of a subset of genes with distal enhancers. Accordingly, the deletion of CTCF sites flanking the *Il12b* gene reduced IL-12 production by cDC1s. Our data reveal an essential role of cohesin-mediated chromatin regulation in cell differentiation and function *in vivo*, and its bi-directional crosstalk with lineage-specifying transcription factors.

## Introduction

A fundamental level of epigenetic regulation in eukaryotic cells is genome organization, the hierarchical folding of chromatin in 3D space that is critical for genome stability and function. High-resolution chromosome capture-based technologies have revealed sub-megabase topologically associating domains (TADs) as a hallmark of higher order chromatin organization (*1, 2*). TADs are formed through the ATP-dependent action of the cohesin complex, which extrudes chromatin loops until it is halted by the protein CTCF bound in the proper orientation (*3, 4*). Cohesin-mediated TAD formation is thought to enhance the fidelity of transcriptional control by facilitating enhancer-promoter interactions, particularly those occurring over long distances (*5–7*). It is also thought to optimize temporal control of gene expression during development (*8*), in part by counteracting Polycomb-mediated repression (*9*). Independent of loop extrusion, the genome is spatially compartmentalized based on the transcriptional and histone modification status of chromatin, with transcriptionally active chromatin located centrally (A compartments) and inactive chromatin located peripherally (B compartments) within the nucleus (*10*).

Acute depletion of cohesin in post-mitotic cells *in vivo* demonstrated its central role in chromosome organization during interphase (*11*). However, because cohesin is strictly required for cell division, understanding the functional consequences of its division-independent functions has been challenging (*12*). The role of cohesin/CTCF-mediated chromatin organization has been demonstrated in special cases such as antigen receptor gene rearrangement in lymphocytes (*13–15*), stochastic protocadherin expression (*16, 17*) or organization of multilobular nuclei (*18*). On the other hand, the effects of cohesin on the terminal differentiation and function of various postmitotic cell types *in vivo* have not been firmly established. In addition to cohesin, various lineage-specifying transcription factors (TFs) have been implicated in the control of 3D genome organization (*19–22*). However, the respective roles of and potential crosstalk between TFs and cohesin in shaping chromatin architecture remain to be explored.

Dendritic cells (DCs) bridge innate and adaptive immunity, linking the recognition of pathogen-derived molecular patterns (PAMPs) to the priming of antigen-specific naive T cells (*23*). The recognition of PAMPs via pattern recognition receptors (PRRs) on DCs leads to DC activation, production of immunostimulatory cytokines and antigen presentation to T cells. DC progenitors in the bone marrow (BM) continuously undergo Flt3 ligand (Flt3L)-dependent differentiation into conventional DCs (cDCs), the primary antigen-presenting DC subset, and plasmacytoid DCs (pDCs), which specialize in type I interferon (IFN-I) production (*24, 25*). BM-derived cDC progenitors exit into the periphery and undergo terminal differentiation driven by local signals such as Notch2 and lymphotoxin (*26–28*). Following their differentiation, all DCs are largely non-proliferative and short-lived with a lifespan of several days. cDCs themselves are composed of two developmentally, transcriptionally and functionally distinct subsets, cDC1s and cDC2s (*29*). A distinct feature of cDC1s is their ability to cross-present exogenous antigens on MHC class I to CD8^+^ T cells (*30*). Furthermore, cDC1s are a critical source of the cytokine interleukin-12 (IL-12) during infection with intracellular pathogens as well as in the tumor microenvironment, which has made them an attractive target in evolving cancer therapies (*31*). Among the multiple TFs that control the differentiation of DC subsets, interferon regulatory factor 8 (IRF8) is induced early in DC specification and controls the development of committed DC progenitors, pDCs and cDC1s (*32–35*).

Here we report that cohesin/CTCF-mediated genome organization enables the post-mitotic differentiation and function of cDCs *in vivo*, including the hallmark functions of cDC1s such as cross-presentation and IL-12 production. Notably, we found that cohesin and IRF8 play complementary roles in establishing DC chromosome organization: while cohesin maintained TADs and related patterns of chromosome structure, IRF8 was essential for proper compartmentalization. The loss of cohesin impaired the induction of select activation-induced genes, including *Il12b* that encodes IL-12. Notably, cohesin-dependent inducible genes were driven by more distal enhancers and were also enriched in Polycomb repression, establishing the role of cohesin in these processes. Furthermore, we found that CTCF sites flanking the *Irf8* locus were required for optimal IRF8 expression, while CTCF sites forming the *Il12b* gene TAD facilitated the IL-12 response by DCs. Our results reveal an essential role of cohesin-mediated chromatin organization in lineage-specific cell differentiation and activity *in vivo*, and suggests mechanisms whereby cohesin drives long-range gene activation and counteracts polycomb repression.

## Results

### Cohesin is required for the differentiation of conventional dendritic cells

To test the role of the cohesin complex in the DC lineage, we targeted its essential subunit Smc3. Unlike other subunits of the cohesin complex, the loss of Smc3 was shown to abolish the binding of the complex to DNA (*36*). We crossed mice with a LoxP-flanked (“floxed”) allele of *Smc3* (*Smc3*^fl^, (*37*)) to the *CD11c-*Cre deleter strain (*38*) to generate mice with DC-specific deletion of *Smc3* (*Smc3*^ΔDC^). Cre recombination in the *CD11c-*Cre strain occurs after the commitment of progenitors to the DC lineage, is highly efficient in mature cDCs and long-lived, tissue-resident CD11c^+^ cells (e.g. Langerhans cells and alveolar macrophages), but is less efficient in pDCs (*38*). Indeed, cDCs isolated from spleens of *Smc3*^ΔDC^ mice harbored excised *Smc3* alleles (Fig. S1A), confirming the robust DC-specific deletion of Smc3. The analysis of splenic DCs from *Smc3*^ΔDC^ mice revealed a ∼50% reduction in the absolute numbers (Fig. 1A) and frequencies (Fig. S1B) of cDC1 and of the Notch2-dependent Esam^+^ subset of cDC2 (*27*). In contrast, Esam^-^ cDC2 and pDCs were unaffected in *Smc3*^ΔDC^ mice (Fig. 1A, S1B,C). We also observed a near-complete absence of cDC1s in the lung (Fig. 1B), and a reduction of skin-derived migratory cDCs in the skin-draining lymph nodes of *Smc3*^ΔDC^ mice (Fig. S1D). Finally, epidermal Langerhans cells and alveolar macrophages were nearly absent from the skin and lungs of *Smc3*^ΔDC^ mice, respectively (Fig. S1E,F). Because these CD11c^+^ populations are established before birth and maintained in tissues by local self-renewal (*39*), their depletion was likely due to the role of cohesin in cell division (*12*). To exclude a similar role in the observed DC phenotype, we measured DC turnover by administering the nucleoside analog EdU for 3 days. In this time frame, proliferating DC progenitors are expected to incorporate EdU and give rise to mature DCs, as evidenced by the detection of EdU in splenic cDCs (Fig. 1C). In the spleens of *Smc3*^ΔDC^ mice, EdU incorporation was marginally reduced in Esam^+^ cDC2s and normal in cDC1s, suggesting a normal cell turnover (Fig. 1C). Thus, the post-commitment loss of Smc3 impaired cDC1 homeostasis independently of proliferation.

**Figure 1.**
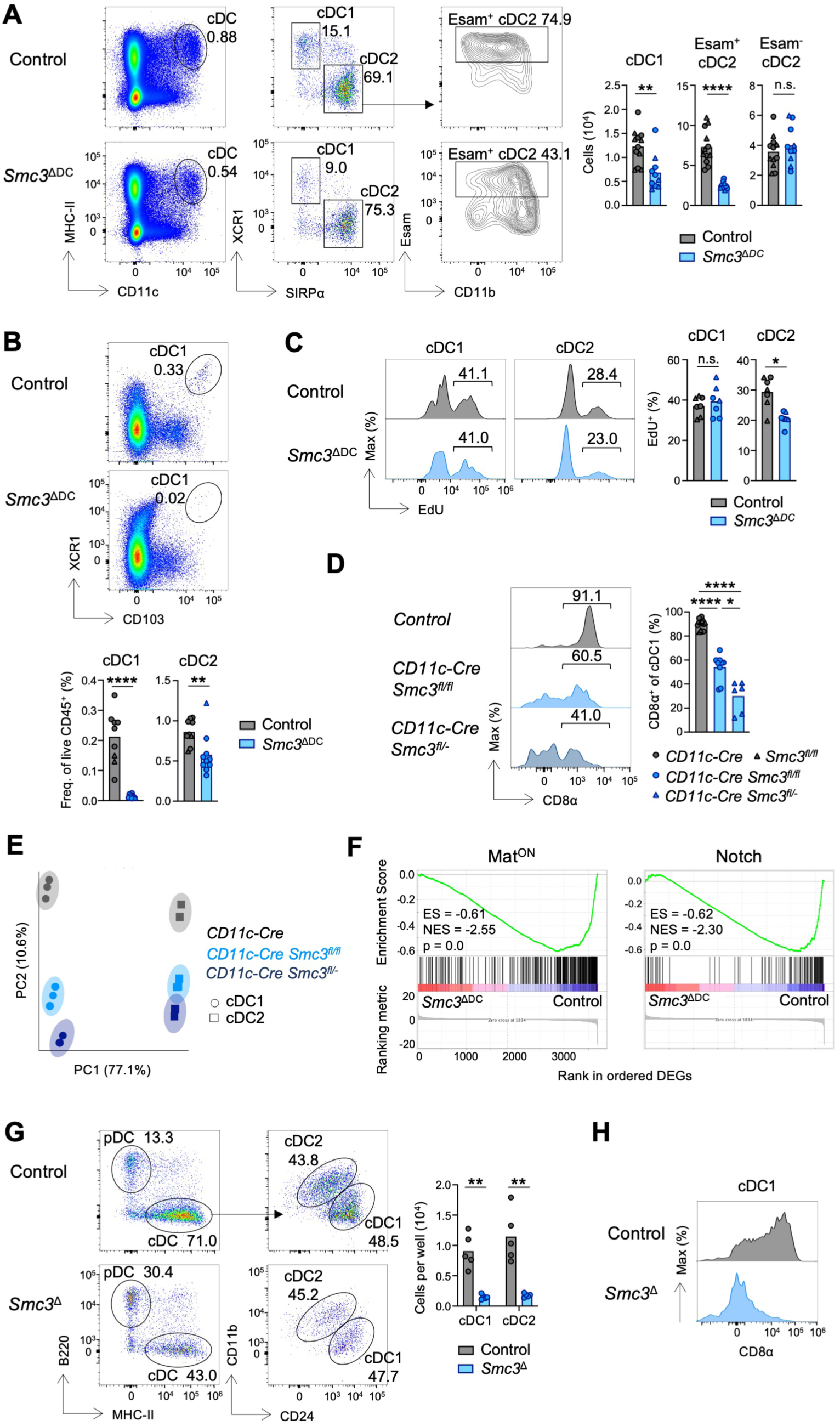
Cohesin is Required for cDC Differentiation Independently of Proliferation. (A) Representative flow plots (left) and numbers (right) of splenic cDC populations. cDC1 = Lin^-^ (TCR-β^-^ CD3^-^ CD19^-^ NKp46^-^ NK1.1^-^) CD11c^+^ MHC-II^+^ XCR1^+^ SIRPα^-^, cDC2 = Lin^-^ CD11c^+^ MHC-II^+^ XCR1^-^ SIRPα^+^. Pooled from two independent experiments. (B) Representative flow plots (top) and frequencies (bottom) of lung cDC populations. Pooled from two independent experiments. (C) Mice were injected on three consecutive days with EdU. Representative histograms (left) and frequencies (right) of EdU^+^ splenic cDCs one day after the last injection. (D) Representative histograms (left) and frequencies (right) of CD8α expression on splenic cDC1s. Pooled from three independent experiments. (E-F) RNA-seq was performed on purified splenic cDCs. Shown are PCA (panel E) and GSEA (panel F) of Mat^ON^ (*35*) and Notch (*23*) signatures among differentially expressed genes (DEGs) between control and *Smc3*^Δ^ cDC1s. ES = enrichment score, NES = normalized enrichment score. (G-H) Bone marrow cells from *R26^CreER/^*^+^ (Control) or *R26^CreER/^*^+^ *Smc3^fl/fl^* (*Smc3*^Δ^) mice were differentiated into DCs *in vitro* in the presence of Flt3L and OP9-DL1 stromal cells for 8 days. 4-hydroxytamoxifen (4-OHT) was added to cultures at day 3 to induce recombination of the floxed *Smc3* allele. Shown are representative flow plots (left) and numbers (right) of indicated cDC subsets (panel G), and representative histograms of CD8α on cDC1s (panel H). Symbols represent individual mice, and bars represent mean. For panels A-D, Control = *CD11c-*Cre (grey circles) and *Smc3^fl/fl^* (grey triangles), *Smc3*^ΔDC^ = *CD11c-*Cre *Smc3^fl/fl^* (blue circles) and *CD11c-*Cre *Smc3^fl/-^* (blue triangles). Statistical differences were evaluated using Mann-Whitney test. *p < 0.05, **p < 0.01, ****p < 0.0001; n.s. not significant.

Flow cytometry analysis of the remaining XCR1^+^ *Smc3*^ΔDC^ cDC1s revealed the loss of the hallmark phenotypic marker CD8α, which was more profound in mice with one germline null allele of *Smc3* (*CD11c*-Cre *Smc3*^fl/-^) (Fig. 1D). Furthermore, *Smc3*^ΔDC^ Esam^+^ cDC2s exhibited aberrant upregulation of the integrin CD11b (Fig. S1G), suggesting that the terminal differentiation of both cDC1s and cDC2s was affected. To test this notion, we performed bulk RNA sequencing of sorted splenic cDC1s and cDC2s from *CD11c*-Cre *Smc3*^fl/fl^, *CD11c*-Cre *Smc3*^fl/-^ and control *CD11c*-Cre animals. By principal component analysis (PCA), PC1 resolved cDC subsets, whereas a cohesin-dependent signature that scaled with cohesin gene dosage drove PC2 (Fig. 1E). Notably, the differences between *Smc3*^Δ^ and control cDCs along PC2 were greater for cDC1s than cDC2s (Fig. 1E). Gene set enrichment analysis (GSEA) revealed an enrichment of the Mat^ON^ signature, a set of genes conserved during cDC1 homeostatic maturation in several tissues (*40*), among those DEGs downregulated in *Smc3*^ΔDC^ cDC1s (Fig. 1F). A similar enrichment was observed for target genes of Notch2, an important signal for terminal cDC maturation (*27, 28*), even after removing genes shared with the Mat^ON^ signature (Fig. 1F, S1H,I). In contrast, proliferation signature genes were not enriched among the DEGs downregulated in *Smc3*^ΔDC^ cDCs, consistent with the low proliferation of cDCs that was unaffected by Smc3 deletion. Collectively, these data reveal a requirement for cohesin in the *in vivo* differentiation of cDCs, particularly cDC1s and Esam^+^ cDC2s.

To confirm the role of cohesin in cDC differentiation *in vitro*, we crossed the *Smc3*^fl^ mice to mice expressing a tamoxifen-inducible Cre recombinase-estrogen receptor fusion protein (CreER) from the ubiquitous *Rosa26* locus (*R26^CreER^*). The development of the major DC lineages (cDC1, cDC2, pDC) can be modeled by culturing total BM cells with Flt3L for 7-8 days (FL-BMDC cultures) (*41*). In FL-BMDC cultures from *R26^CreER^ Smc3*^fl/fl^ mice, the addition of 4-hydroxytamoxifen (4-OHT) at day 3 of culture, after the peak of progenitor proliferation has occurred, permitted DC development but substantially depleted Smc3 protein in most cells by day 8 (Fig. S2A-C). Notably, such Smc3-deleted (*Smc3*^Δ^) FL-BMDC cultures showed a significant reduction of pDCs (∼1.5-fold), cDC2s (∼2-fold) and especially cDC1s (∼4-fold) compared to cultures from control *R26^CreER^* mice (Fig. S2D). We purified the residual *Smc3*^Δ^ DC subsets from FL-BMDC cultures and analyzed them by RNA-seq. By PCA, control and *Smc3*^Δ^ pDCs and cDC2s clustered together, whereas *Smc3*^Δ^ cDC1s clustered nearest to control cDC2s (Fig. S2E). Furthermore, DEGs upregulated in *Smc3*^Δ^ cDC1s demonstrated enrichment of the signature of primary splenic cDC2s (*42*), suggesting that cohesin optimizes cDC1 subset specification in this system (Fig. S2F). The induction of Notch2 signaling, via the co-culture with OP9 stromal cells expressing the Notch ligand Delta-like 1 (OP9-DL1), decreases cell yield but facilitates the differentiation of *bona fide* cDC1s and Esam^+^ cDC2s (*43*). In these conditions, the output of both cDC1s and cDC2s was dramatically reduced by Smc3 deletion (Fig. 1G), and *Smc3*^Δ^ cDC1s failed to upregulate CD8α (Fig. 1H). These data confirm the function of cohesin in cDC differentiation, particularly in the context of physiological signals such as Notch ligands.

### Cohesin controls the function of conventional dendritic cells in vivo

The impaired differentiation of cohesin-deficient cDCs prompted us to test their function, including responses to PAMPs mediated via their cognate PRRs such as Toll-like receptors (TLRs). RNA-Seq of *Smc3*^Δ^ cDCs *ex vivo* (Fig. 1E) and from cultures (Fig. S2E) showed reduced expression of *Il12b*, the gene encoding the main p40 isoform of IL-12 (Fig. 2A). Accordingly, *Smc3*^Δ^ FL-BMDC showed significantly less IL-12p40 production when untreated or treated with the TLR9 ligand unmethylated CpG oligonucleotides (CpG) or with the TLR11 ligand profilin (Fig. 2B). During acute infection with *Toxoplasma gondii* or its extract soluble tachyzoite antigen (STAg) containing profilin, cDC1s are the critical source of IL-12 required for host resistance (*44–46*). We found that *Smc3*^ΔDC^ mice showed a dramatic loss of serum IL-12p40 in the steady state (Fig. 2C). The challenge with profilin elicited robust production of both IL-12p40 and IL-12p70 in control mice, yet this response was completely ablated in *Smc3*^ΔDC^ mice (Fig. 2C,D). After infection with *Toxoplasma gondii*, *Smc3*^ΔDC^ mice showed reduced serum IL-12p40 and reduced production of IL-12p40 by peritoneal exudate cells and splenocytes (Fig. 2E). Thus, cohesin is required for both homeostatic and PAMP-induced production of IL-12 by cDC1s.

**Figure 2.**
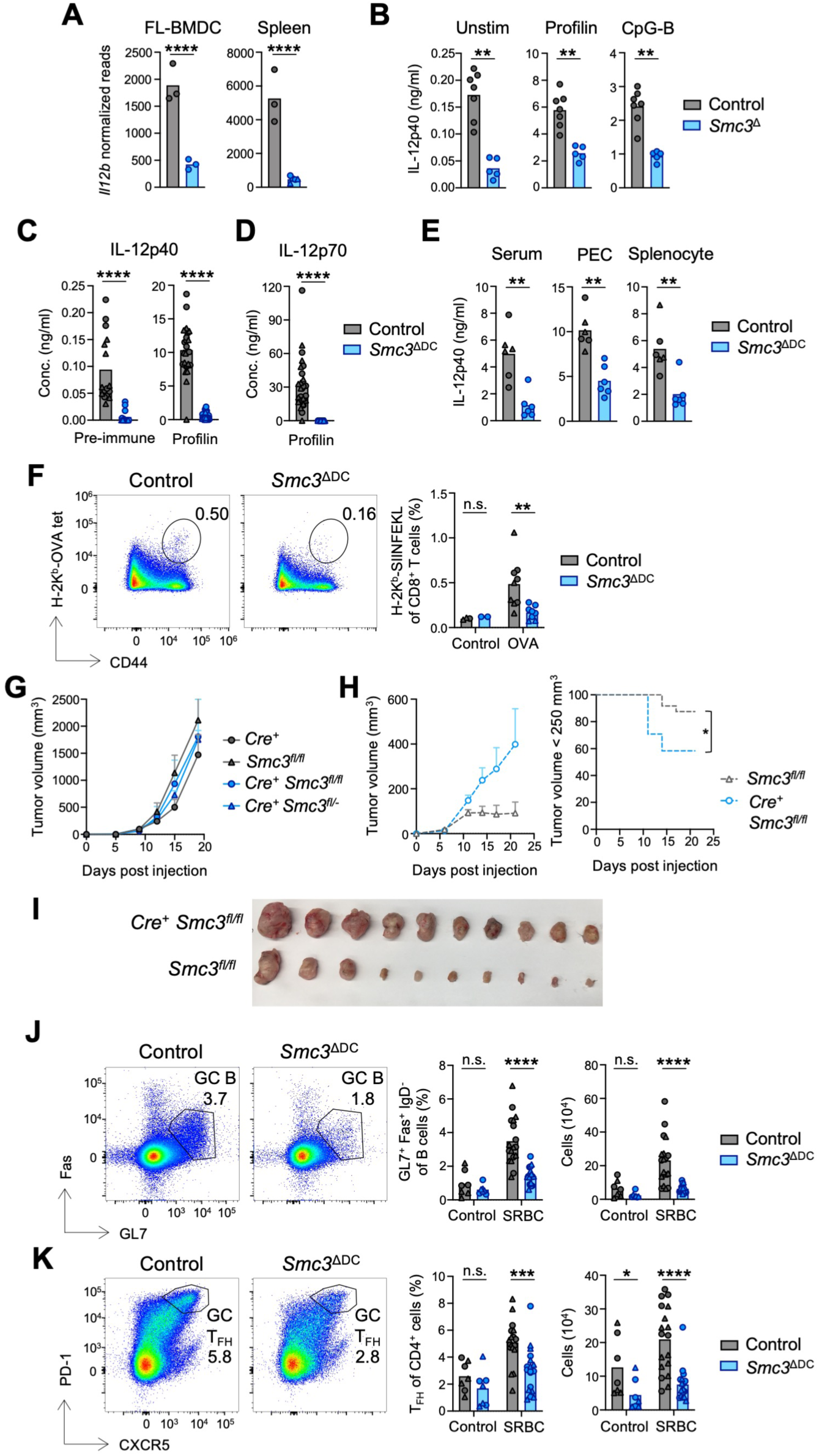
Cohesin is Required for cDC Function in vivo. (A) Normalized read counts for *Il12b* gene from RNA-Seq of cDC1s purified from FL-BMDC cultures (left) and spleen (right). (B) FL-BMDCs were stimulated with profilin (middle), CpG-B (right) or left unstimulated as a control (left). Quantification of IL-12p40 concentration in supernatant of indicated cultures 24 hours later. (C-D) Quantification of IL-12p40 (panel C) and IL-12p70 (panel D) concentration in the serum of indicated mice both prior to and 6 hours post challenge with profilin. Pooled from two independent experiments. (E) Quantification of IL-12p40 in the serum (left) and from peritoneal exudate cell (PEC) (middle) and splenocyte (right) supernatant 5 days after infection with *T. gondii*. (F) Indicated mice received either OVA-pulsed or control BALB/c splenocytes. Shown are representative flow plots (left) and fractions (right) of endogenous OVA-specific CD8^+^ T cells in spleen on day 8 post-transfer. (G) Mice received bilateral subcutaneous implantation of MC38 tumor cells. Shown are tumor growth curves in indicated mice. (H-I) Mice received bilateral subcutaneous implantation of MC38 tumor cells and were treated with immunotherapy (αPD-1 + αCD137) every third day beginning day 6 post-tumor inoculation. Shown are tumor growth curves (panel H, left), Kaplan-Meier curves showing mice in which tumor burden exceeded 250 mm^3^ (panel H, right), and largest volume tumors from each group at day 21 experiment endpoint (panel I). (J-K) Mice were injected either with sheep RBCs or PBS, and the germinal center response was analyzed in the spleen on day 8. Shown are representative flow plots (left), frequencies (middle) and numbers (right) of germinal center B cells (B220^+^ GL7^+^ Fas^+^ IgD^-^) (panel J) and T_FH_ cells (TCR-β^+^ CD4^+^ PD-1^+^ CXCR5^+^) (panel K). Pooled from two independent experiments. In bar plots, symbols represent individual mice, and bars represent mean. In panels A-F, J-K, Control = *CD11c-*Cre (grey circles) and *Smc3^fl/fl^* (grey triangles), *Smc3*^ΔDC^ = *CD11c-* Cre *Smc3^fl/fl^* (blue circles) and *CD11c-*Cre *Smc3^fl/-^* (blue triangles). In tumor experiments, symbols represent the mean of all tumors (two per mouse). Statistical differences were evaluated using Mann-Whitney test, except for the log-rank (Mantel-Cox) test for evaluating the Kaplan-Meier curve (H). *p < 0.05, **p < 0.01, ***p < 0.001, ****p < 0.0001; n.s. not significant.

To test the ability of cohesin-deficient cDC1s to cross-present cell-associated antigens, we injected mice with allogeneic (H-2K^d^) splenocytes loaded with ovalbumin (OVA) protein and measured by tetramer staining the expansion of OVA-specific CD8^+^ T cells, a response fully dependent on endogenous cDC1s (*42*). Whereas control mice were capable of cross-priming CD8^+^ T cells specific to OVA, *Smc3*^ΔDC^ mice were defective in this capacity (Fig. 2F). To determine whether cohesin-dependent cDC1 cross-presentation was important in the context of anti-tumor immunity, we implanted MC38 adenocarcinoma cells in both flanks of control and *Smc3*^ΔDC^ mice. Six days later, we initiated an immunotherapy regimen consisting of PD-1 blockade (αPD-1) and agonist antibody to CD137 (4-1BB), a combination that requires cDC1s to elicit a productive anti-tumor CD8^+^ T cell response (*47*). Without immunotherapy, MC38 tumors grew at comparable rates in all control (*CD11c*-Cre or *Smc3*^fl/fl^) and *Smc3*^ΔDC^ (*CD11c*-Cre *Smc3*^fl/fl^ or *CD11c*-Cre *Smc3*^fl/-^) mice (Fig. 2G). Immunotherapy impaired tumor growth in control but not in *Smc3*^ΔDC^ mice (Fig. 2H), which also showed a significant increase in larger tumors (Fig. 2H,I). Thus, cohesin regulates cross-presentation by cDC1s and the resulting cDC1-dependent antitumor response.

cDC2s are specialized in antigen presentation to CD4^+^ T cells, and Esam^+^ cDC2s in particular mediate the development of follicular helper T (T_FH_) cell differentiation and germinal center (GC) reactions after immunization (*48*). We therefore immunized control and *Smc3*^ΔDC^ mice with sheep red blood cells (SRBCs) and analyzed the development of GC T_FH_ and B cell responses. In line with the diminished Esam^+^ cDC2 population, *Smc3*^ΔDC^ mice showed a significant reduction of SRBC-elicited GC B cells (Fig. 2J) and T_FH_ cells (Fig. 2K). Collectively, these data demonstrate that cohesin is required for cDCs to acquire and execute their hallmark functional features *in vivo*.

### Cohesin-mediated genome organization in DCs

Given the observed proliferation-independent role of cohesin in cDC differentiation and function, we directly tested its effect on the genome organization of DCs. To this end, we purified cDCs (both subsets) and pDCs from 4-OHT-treated control and *Smc3*^ΔDC^ FL-BMDC cultures (Fig. S2) and performed genome-wide chromosome conformation capture (Hi-C) (Fig. 3A). Pairwise comparison of Hi-C maps indicates global similarities and differences between chromosome folding in studied cells. PCA of pairwise Hi-C map similarity revealed the separation of both control and *Smc3*^Δ^ cDCs and pDCs along PC1 (Fig. S3A), suggesting that cohesin depletion does not erase the divergence between DC subsets. Nevertheless, pDCs and especially cDCs showed a clear separation between control and Smc3-deficient cells along PC2, indicating major changes in chromosome organization upon cohesin depletion.

**Figure 3.**
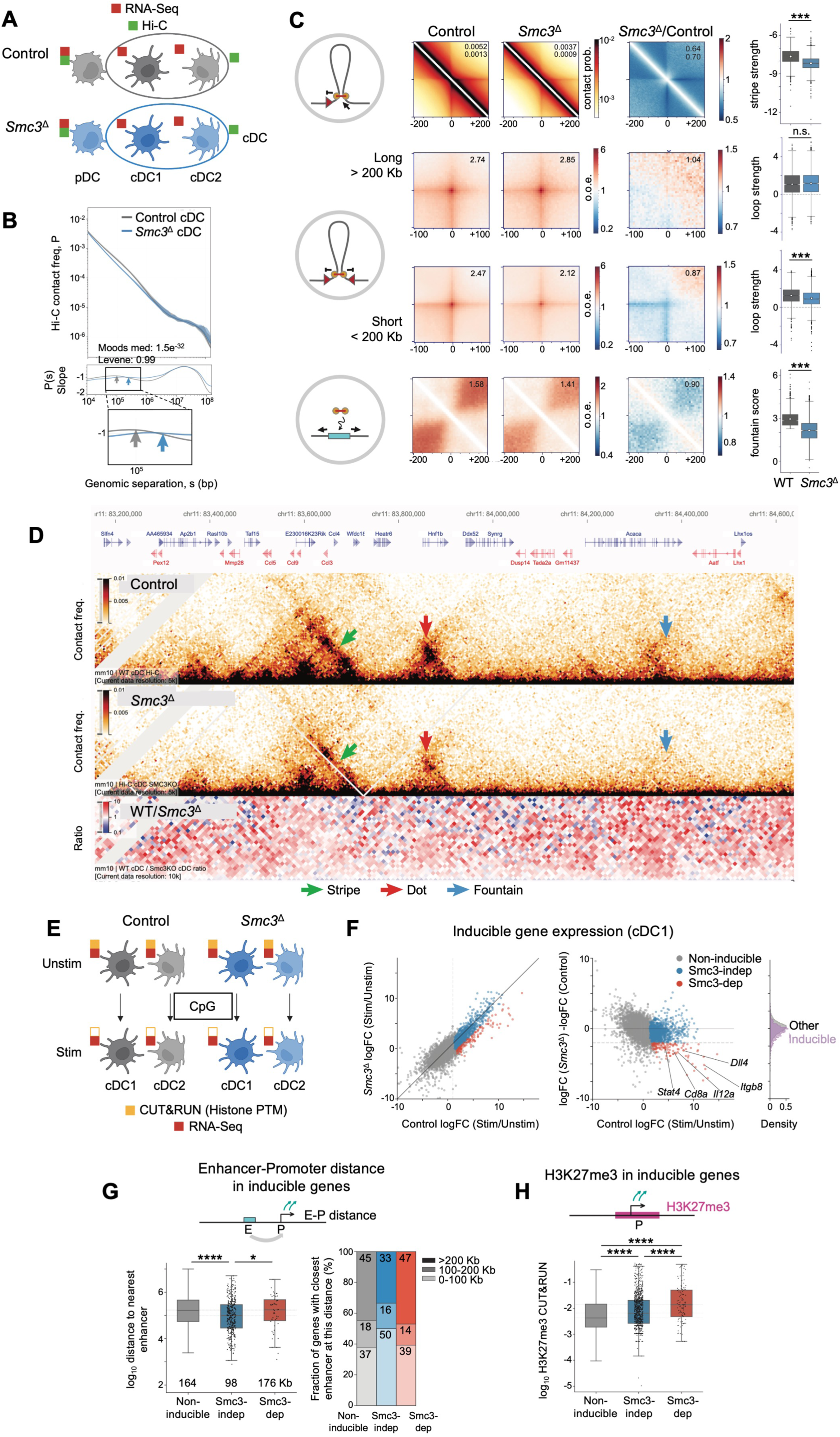
Cohesin Regulates Genome Organization in DCs. (A) Experimental schematic. FL-BMDCs cultures were established from *R26^CreER/^*^+^ (Control) or *R26^CreER/^*^+^ *Smc3^fl/fl^* (*Smc3*^Δ^) mice, treated with 4-OHT on day 3, and on day 8 analyzed by RNA-Seq for individual DC subsets, or by Hi-C for cDCs and pDCs. (B) P(s) curves (top) and their slopes (bottom) for cDCs. Shaded regions represent the 90% confidence interval at each genomic separation range. Statistical differences between the distributions of slopes at 50-150 Kb were calculated with Moods median test and Levene test for variance equivalence. (C) Average pileups around forward-oriented CTCF peaks (stripes, top row), around pairs of both short-range or long-range CTCF peaks with convergent orientation (dots, middle two rows) and at active chromatin (fountains, bottom row) (left) and summary statistics of their strength (right) in cDCs from indicated samples. (D) Contact frequency maps at a representative locus highlighting the above-mentioned chromatin structures in *Smc3*^Δ^ and control cDCs. (E) Experimental schematic. Control and *Smc3*^Δ^ FL-BMDCs were either left unstimulated or stimulated for 6 hours with CpG. cDC1s and cDC2s were purified and their transcriptomes analyzed by RNA-seq and (for unstimulated cells only) by CUT&RUN for histone modifications. (F) Scatterplots of genes in control and *Smc3*^Δ^ cDC1s with or without CpG activation. Dashed lines (right) represent the selection criteria for the gene categories. “Smc3-dependent genes” were defined as those induced by CpG in control cDC1s (log_2_FC > 2, p < 0.01), but not to the same extent in *Smc3*^Δ^ cDC1s (WT log_2_FC / *Smc3*^Δ^ log_2_FC > 2). “Smc3-independent genes” were defined as those that satisfy only the first condition. “Non-inducible genes” were not significantly upregulated in control cDC1s upon stimulation. Select Smc3-dependent genes are indicated. (G) Distance of Smc3-dependent, Smc3-independent and non-inducible genes to their nearest enhancer in cDC1s (left). Dashed lines represent the medians of the distributions, and the median value in Kb is indicated on the plot. Statistical differences were calculated with the Mann-Whitney one-sided test. Proportion of indicated gene subsets that have their nearest enhancer located within 100 Kb, 100-200 Kb or more than 200 Kb away from their transcription start site (right). (H) Magnitude of CUT&RUN-derived H3K27me3 in cDCs within the insulated neighborhood of genes from the indicated gene categories. Dashed lines represent the medians of the distributions. Statistical differences were calculated with the Mann-Whitney one-sided test. Data are presented as box plots. *p < 0.05, ***p < 0.001, ****p < 0.0001; n.s. not significant.

We examined global changes in Hi-C maps upon cohesin depletion. The curves of the contact frequency P(s) as a function of genomic separation (s) are used to detect extruded loops, seen as a characteristic hump (elevated contact frequency) at the scale corresponding to the loop size (*49, 50*). While this curve for control DCs showed a hump corresponding to ∼100 Kb loops, this hump was significantly reduced and the curve flattened in *Smc3*^Δ^ cDCs (Fig. 3B) and *Smc3*^Δ^ pDCs (Fig. S3B). To estimate the degree of cohesin depletion in DCs, we used Hi-C data from mouse embryonic stem cells with degron-mediated depletion of cohesin (Fig. S3C). Using *in silico* mixtures containing different fractions of residual cohesin (*51*) suggested ∼54% and ∼73% of residual cohesin activity in *Smc3*^Δ^ cDCs and pDCs compared to their respective wild-type levels (Fig. S3D,E). These estimates suggest that the profound cohesin depletion in *Smc3*^Δ^ FL-BMDC cultures (Fig. S2B,C) translates into a ∼46% loss of cohesin function in cDCs.

While the overall compartmentalization of the genome as defined by the first eigenvector (E1) was unchanged in *Smc3*^Δ^ cDCs (Fig. S3F), the latter displayed weakened insulation at CTCF sites, revealing the expected dissolution of TADs (Fig. S3G). Moreover, *Smc3*^Δ^ cDCs manifested the weakening and shortening of “stripes” emanating from CTCF sites (Fig. 3C,D). Such stripes reflect the process of extrusion by cohesin that was stopped at a CTCF site, with many stripes associated with promoters, enabling enhancer-promoter interactions (*10*). We also observed the weakening of “dots”, which reflect the stopping of cohesin at two CTCF sites and a transient loop between them (*51*), particularly for the loops <200 Kb (Fig. 3C,D). Finally, we examined the recently described cohesin-associated features known as “jets” or “fountains”, which reflect the loading of cohesin at active enhancers (*52, 53*). We identified an average “fountain” by aggregating Hi-C maps at active chromatin that shows such a pattern, and observed a significantly diminished fountain signature in the *Smc3*^Δ^ cDCs (Fig. 3C,D). Similar effects, albeit to a lower degree, were observed in *Smc3*^Δ^ pDCs (Fig. S3H). Altogether, cohesin-deficient cDCs manifest genome-wide reductions in major features associated with loop extrusion.

### Cohesin regulates a subset of the cDC activation program

Acute depletion of cohesin impaired the expression of inducible genes in TLR-stimulated macrophages (*54*). To dissect the role of cohesin in activation-induced DC responses, we stimulated *Smc3*^Δ^ and control FL-BMDC cultures (Fig. S2) with CpG (both A- and B-types), which activate all DC subsets through TLR9. After 6 hours, unstimulated or CpG-stimulated cDC1 and cDC2 subsets were sorted and analyzed by RNA-seq (Fig. 3E). PCA of RNA-seq profiles showed that both control and *Smc3*^Δ^ cDCs separated equally from their unstimulated counterparts along PC1 (Fig. S3I), suggesting that the overall activation of cohesin-deficient DCs was not grossly impaired. Control and *Smc3*^Δ^ DCs separated along PC2, with activated DCs showing larger divergence, revealing an added effect of cohesin during activation. Indeed, a small subset of activation-induced genes (e.g. *Shisa3*, *Cxcl11*, *Il6* and *Saa3*) showed preferential expression in both subsets of control cDCs. In addition, several genes showed higher expression in control activated cDC1s, including important inflammatory mediators *Il12b* and *Nos2* (Fig. S3J). Thus, the loss of cohesin does not block the cDC activation program but impairs the induction of a specific subset of this program.

Consistent with PCA, pairwise comparison between control and *Smc3*^Δ^ cDCs showed only a slight overall reduction of activation-induced genes in the latter, except for a small subset of activation-induced genes that were strongly reduced (Fig. 3F and S4A). To analyze cohesin-dependent inducible genes, we defined them as those induced by CpG in control but not *Smc3*^Δ^ DCs (n=117 and 112 in cDC1s and cDC2s, respectively), versus Smc3-independent genes that were induced in both control and *Smc3*^Δ^ DCs (n=989 and 1348 in cDC1s and cDC2s, respectively), and non-inducible genes that were not significantly upregulated upon stimulation (n=7501 and 7186 in cDC1s and cDC2s, respectively) (Fig. 3F and S4A). We then used CUT&RUN to profile histone modifications in wild-type non-activated cDC1s and cDC2s (Fig. 3E), defined active enhancers for all transcribed genes and determined the distances between the promoter to the nearest enhancer for each gene. We observed that Smc3-dependent inducible genes were >1.7-fold more distant from the nearest enhancer than Smc3-independent inducible genes (Fig. 3G and S4B). Moreover, 47.1% of Smc3-dependent inducible genes in cDC1 were >200 Kb from the nearest enhancer in comparison to about 33.2% for Smc3-independent genes (Fig. 3G) (40.2% and 19.3%, respectively, in cDC2s, Fig. S4B).

We further compared the three groups of genes using histone profiling described above, as well as ATAC-Seq and ChIP-Seq for CTCF from *in vitro*-derived cDCs described below in Fig. 4. Consistent with their increased distance to the enhancer as shown above, Smc3-dependent genes showed a significantly increased distance to the nearest CTCF binding site compared to Smc3-independent genes (Fig. S4C). They also showed a significant reduction of open chromatin peaks as defined by ATAC-Seq (Fig. S4D). Among histone modifications, the most striking difference was in the histone H3 tri-methylation at lysine 27 (H3K27me3), which was significantly enriched in Smc3-dependent vs Smc3-independent genes, and generally in inducible compared to non-inducible genes in both cDC1 and cDC2 (Fig. 3H and S4E). H3K27me3 is a mark of repression by the Polycomb repressive complex, consistent with the role of cohesin in disrupting Polycomb-dependent chromatin interactions (*9*). Other significant differences between Smc3-dependent vs -independent genes included increased histone H3 tri-methylation at lysine 9 (H3K9me3) in both cDC1 and cDC2 (Fig. S4F), a mark generally associated with heterochromatin. They also included decreased histone H3 tri-methylation at lysine 4 (H3K4me3) in cDC1 only (Fig. S4G), a mark associated with active gene promoters. Both of these differences, along with reduced ATAC-Seq signal, are consistent with increased Polycomb mark on Smc3-dependent genes. No differences were observed in H3 lysine 4 monomethylation or lysine 27 acetylation (Fig. S4H,I). Collectively, these data suggest that cohesin in DCs is required for the induction of a distinct group of genes that are located farther from their enhancers and CTCF-binding sites, and are subject to Polycomb-mediated repression.

**Figure 4.**
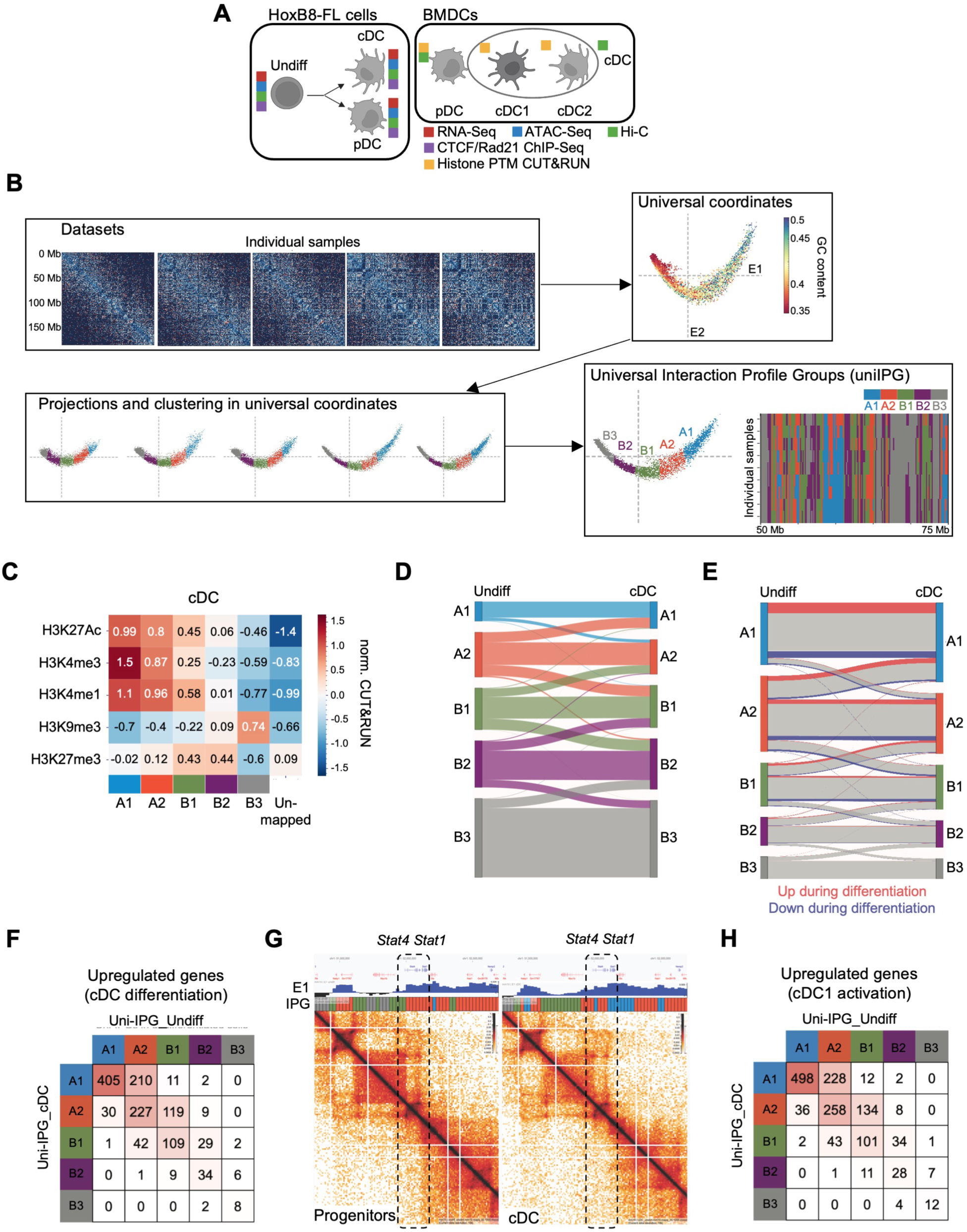
Uni-IPG Analysis Reveals Dynamics of Genome Organization during DC Differentiation. (A) Experimental schematic. RNA-Seq, ATAC-Seq, Hi-C and ChIP-Seq for CTCF and Rad21 were performed on HoxB8-FL progenitors (Undiff) and differentiated cDCs and pDCs, accompanying the datasets performed on FL-BMDCs in Figure 3. (B) Workflow to generate unified interaction profile groups (uni-IPGs). The simultaneous decomposition of all Hi-C subsets was performed on a consensus compartmental profile, which allowed for harmonization across samples and experimental systems. (C) Heatmap of mean enrichment of CUT&RUN-derived histone marks (Z-score normalized log10 signal in 25 Kb bins) in uni-IPG states in HoxB8-FL-derived cDCs. (D) Sankey plot of IPG transitions of genomic bins between progenitors and cDCs. (E) Sankey plot of IPG transitions of genes between progenitors and cDCs for expressed genes, with the fractions of upregulated (red) and downregulated (blue) genes in each compartment indicated. (F) The matrix showing numbers of genes in each uni-IPG in progenitors and cDCs for the genes that were upregulated during cDC differentiation. (G) Hi-C maps in the vicinity of the activation-inducible *Stat4/Stat1* locus in HoxB8-FL progenitors and cDCs. First projected spectral component (E1) and uni-IPG tracks are shown above the Hi-C maps. (H) The matrix showing numbers of genes in each uni-IPG in progenitors and cDCs for the genes that were upregulated during the activation of cDC1s by CpG.

### The role of genome compartmentalization in DC differentiation and activation

A recent analysis of Hi-C data from cell lines revealed additional depth of genome compartmentalization into multiple active and inactive compartments with distinct properties, termed the interaction profile groups (IPG) (*55*). We sought to apply the IPG approach to the dynamics of genome compartments during cell differentiation and activation, using DCs as a paradigm. To compare DCs to their progenitors, we used the Flt3L-dependent HoxB8-FL cells that can be grown as multipotent progenitors or induced to differentiate into a mixture of pDCs and immature cDC2-like cDCs (*56, 57*). We performed Hi-C on HoxB8-FL progenitors and their DC progeny, as well as epigenetic profiling of HoxB8-FL progenitors to match that of BM-derived DCs (Fig. 4A). The resulting Hi-C profiles corresponded to the differentiation trajectory of *ex vivo* DC progenitors (*58*) as determined by PCA (Fig. S5A), with the higher Hi-C resolution and direct progenitor-progeny relationship of the HoxB8-FL system facilitating the analysis of genome compartments. First, we sought to integrate Hi-C profiles of HoxB8-FL progenitors, HoxB8-FL-derived pDCs and cDCs, and BM-derived DCs (Fig. 4A). The standard method of Hi-C decomposition (*59*) generated vastly different compartmental profiles of progenitors and DCs (Fig. S5B). We therefore developed a method for a simultaneous decomposition of all Hi-C subsets and their projection on a consensus compartmental profile, which allowed efficient harmonization of progenitor and DC subsets from both experimental systems (Fig. 4B and S5B). Subsequent clustering of projections yielded two active (A1-A2) and three inactive (B1-B3) unified IPG (uni-IPG) (Fig. 4B), whose identity was consistent with their epigenetic profiles (Fig. 4C). Thus, A1 was enriched in H3K4me3 compared to A2; B1 was enriched both in active chromatin marks and in the Polycomb-associated repressive H3K27me3 mark; B2 was predominantly enriched for H3K27me3; and B3 was enriched in the heterochromatic H3K9me3 mark (Fig. 4C). The analysis of uni-IPG in Smc3-deficient cDCs showed very few differences from controls (Fig. S5C), consistent with cohesin being dispensable for chromatin compartmentalization (*11*).

Progenitor differentiation into cDCs was accompanied by extensive reciprocal transitions between uni-IPG, only a small fraction of which represented “generic” A<>B compartment transitions (Fig. 4D). Most of the transitions were between “adjacent” uni-IPG (e.g. B2<>B1), although “jumps” across an uni-IPG (e.g. B2<>A2) could be observed. Transitions into a more active uni-IPG during differentiation (e.g. A2->A1 or B1->A2) were enriched for upregulated genes, and vice versa (Fig. 4E and S5D). The correlation between uni-IPG transition and the corresponding expression change was particularly striking in “jumps” across one or two uni-IPG (Fig. S5D). Conversely, genes that were upregulated in cDCs vs progenitors either remained in the same uni-IPG or transitioned to a more active uni-IPG (e.g. A2->A1 or B1->A2) (Fig. 4F). Thus, uni-IPG allow a high-resolution analysis of chromatin dynamics in the physiological system of DC differentiation.

We then examined the compartmentalization of genes that were induced by activation in cDCs, as exemplified by the *Stat4/Stat1* locus (Fig. 4G). We observed that the vast majority of them resided in active uni-IPG (A1 or A2) in differentiated cDCs even prior to activation (Fig. 4H and S5E,F). Most of these genes were in the same active uni-IPG in progenitors, whereas others transitioned to a more active compartment (A2->A1 or B1->A2) during differentiation (Fig. 4H). The transitions between compartments also correlated with changes in histone modifications (Fig. S5G). Finally, we examined the relationship between uni-IPG partitioning and cohesin dependence of inducible genes. Consistent with the enrichment of H3K27me3 in cohesin-dependent genes, a larger fraction of them were in B2 or B1 compartments (for cDC1 and cDC2, respectively), comprising a larger net fraction of the H3K27me3-marked B1/B2 compartment (Fig. S5H). Collectively, these data elucidate key properties of the activation-inducible gene expression program in DCs: i) its dependence on cohesin correlates with a larger distance from the enhancer and with the Polycomb-repressed state, and ii) its translocation into the active compartments occurs during differentiation and prior to activation.

### IRF8 controls genome compartmentalization during DC differentiation

Given the extent and relevance of compartment dynamics in DCs, and its apparent independence of cohesin, we hypothesized that it might be controlled by a key DC-specifying TF such as IRF8. To test this notion, we deleted IRF8 from HoxB8-FL cells using CRISPR/Cas9-mediated gene targeting, and characterized the differentiation of the resulting IRF8-deficient (*Irf8*^Δ^) HoxB8-FL cells versus control IRF8-sufficient Hoxb8-FL cells (Fig. 5A). The targeting of *Irf8* resulted in a near-complete loss of IRF8 protein from differentiated HoxB8-FL-derived cells (Fig. S6A). Accordingly, these cells failed to differentiate into pDCs and instead gave rise to immature cDC-like cells lacking MHC class II expression (Fig. 5B). We analyzed the transcriptome of *Irf8*^Δ^ CD11c^+^ DCs by RNA-Seq and integrated it with RNA-Seq of HoxB8-FL cells at different stages of differentiation ((*57*) and this study). By PCA, IRF8-deficient cells mapped onto the normal differentiation trajectory close to immature DCs between differentiation days 2 and 4 (Fig. 5C). Clustering analysis further confirmed the similarity of *Irf8*^Δ^ Hoxb8-FL-derived cells and DCs on day 4 of differentiation, prior to the strong upregulation of subset-specific expression programs (Fig. S6B). Thus, similar to its key role *in vivo*, IRF8 is required for terminal DC differentiation in the HoxB8-FL system.

**Figure 5.**
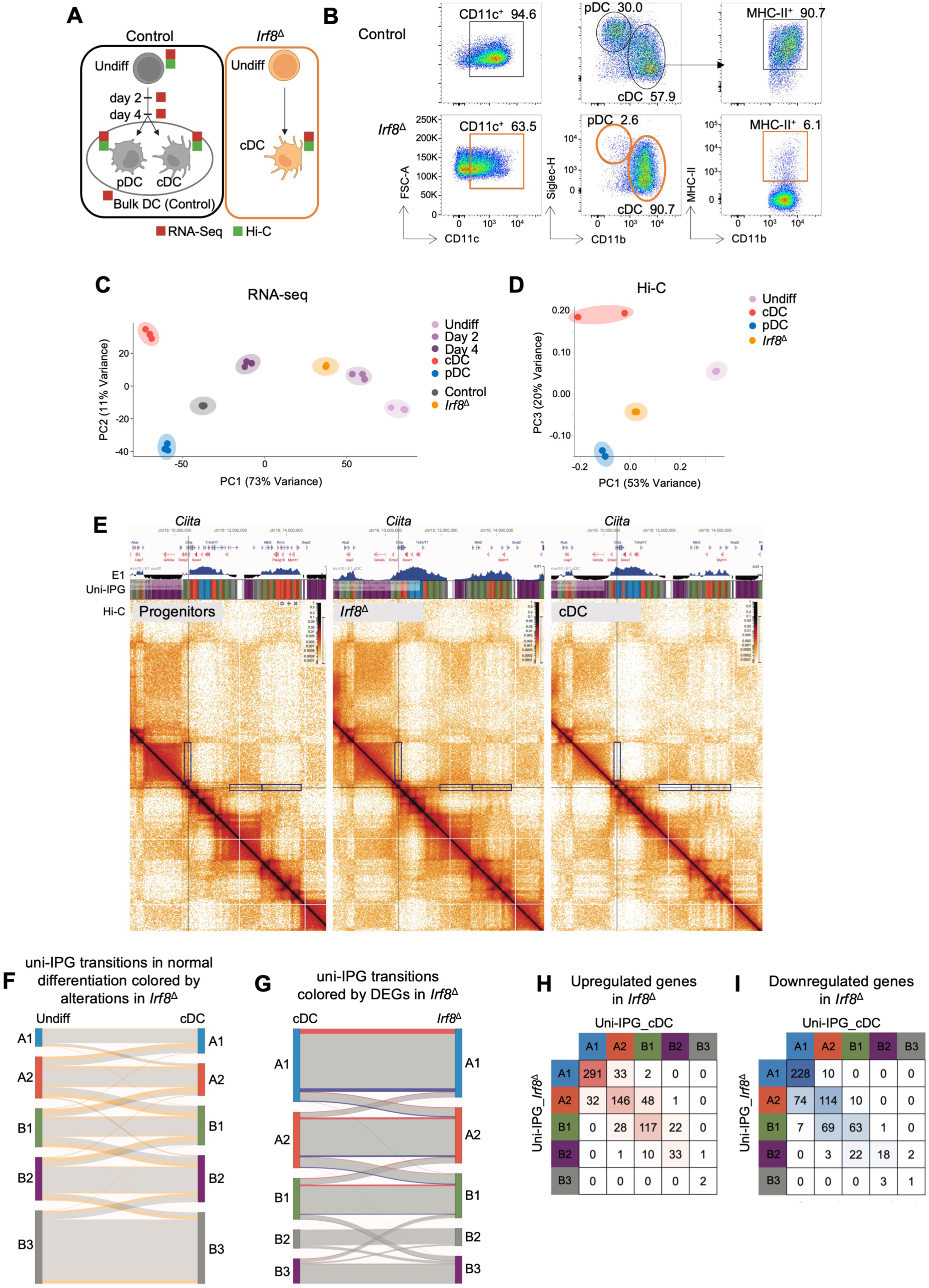
IRF8 Regulates the Genome Compartmentalization of Differentiating DCs. (A) Experimental schematic. RNA-Seq was performed on HoxB8-FL cells at several stages along their differentiation trajectory from undifferentiated progenitors (Undiff) to mature DCs (left). An sgRNA targeting *Irf8* was used to generate IRF8-deficient (*Irf8*^Δ^) HoxB8-FL cells using Cas9-RNPs, and RNA-Seq and Hi-C performed on differentiated *Irf8*^Δ^ DCs as well as RNA-Seq of wild-type control bulk DCs. (B) Representative flow plots of CD11c^+^ cells (left), DC subsets (middle) and MHC-II^+^ cDCs (right). Representative of three independent experiments. (C-D) PCA of RNA-Seq (panel C) and Hi-C (panel D) showing position of *Irf8*^Δ^ DCs relative to the normal transcriptional and chromatin trajectory of differentiating DCs. (E) Hi-C maps in the vicinity of the *Ciita* locus in HoxB8-FL progenitors as well as cDCs and *Irf8*^Δ^ DCs. First projected spectral component (E1) and uni-IPG tracks are shown above the Hi-C maps. Boxed regions highlight compartments with disrupted dynamics in *Irf8*^Δ^ DCs. (F) Sankey plot of IPG transitions of genomic bins during normal differentiation of progenitors into cDCs, colored (in orange) by transitions altered in *Irf8*^Δ^ DCs. (G) Sankey plot of IPG states of genes between cDCs and *Irf8*^Δ^ DCs for expressed genes, with the fractions of upregulated (red) and downregulated (blue) genes in each compartment indicated. (H-I) Matrices showing numbers of upregulated (panel H) and downregulated (panel I) genes in *Irf8*^Δ^ DCs, organized by their IPG status in cDCs and *Irf8*^Δ^ DCs.

We performed Hi-C on IRF8-deficient HoxB8-FL-derived DCs and compared it to Hi-C profiles of wild-type HoxB8-FL progenitors and DCs (Fig. 5A). PCA showed that *Irf8*^Δ^ cells incompletely progressed along the DC differentiation trajectory (PC1) and remained unaffiliated with either pDC or cDC subsets (Fig. 5D). We then compared cohesin-dependent features such as Hi-C contact frequency and the strength of stripes, dots and fountains between undifferentiated progenitors and differentiated wild-type or IRF8-deficient DCs. All these features were different between DCs and progenitors; in contrast, IRF8-deficient cells were either similar to progenitors (for contact frequency, dots and stripes) or half-way between progenitors and DCs (for fountains) (Fig. S6C-F). Thus, IRF8 facilitates optimal cohesin-mediated chromatin organization in DCs.

We then analyzed the compartmentalization of *Irf8*^Δ^ cells using the uni-IPG approach. Representative examples of genes that are upregulated (*Ciita*, Fig. 5E) or downregulated (*Cd34*, Fig. S6G) during DC differentiation showed disrupted compartments in *Irf8*^Δ^ cells. Accordingly, we found that many transitions between uni-IPG in progenitors vs DCs were altered, including the majority of “jumps” over one or two uni-IPG (Fig. 5F and S6H,I). Direct comparison between wild-type cDCs and *Irf8*^Δ^ cells revealed major differences in their uni-IPG structure, which correlated with their gene expression profiles (Fig. 5G and S6J). Indeed, many genes that were up- or downregulated in *Irf8*^Δ^ cells were also located in a different compartment than in wild-type cDCs (Fig. 5H,I). Collectively, these results show that IRF8 controls processes that establish the genome organization and transcriptional profile of differentiated DCs through two mechanisms: by facilitating cohesin-mediated features, and by enforcing cohesin-independent compartmentalization.

### Local genome organization optimizes IRF8 expression in DCs

Having established the role of IRF8 in the control of chromatin dynamics, we asked the opposite question, namely: how does chromatin affect the expression of *Irf8* itself? We reasoned that as an upstream regulator, the *Irf8* locus may have a distinct chromatin arrangement that ensures its proper expression. Indeed, we observed that *Irf8* is located as a sole gene within a compact TAD present in both progenitors and mature DCs (Fig. 6A). This “private” *Irf8* TAD was demarcated by two TAD boundaries (TB1 and TB2), each containing at least three CTCF binding sites (Fig. 6B). Notably, all enhancers that regulate *Irf8* expression in DCs (*60, 61*) and have been shown to physically interact (*62*) are located within this TAD, suggesting that its structure may facilitate *Irf8* expression. To test this notion, we generated mice lacking TB1 and TB2 (*Irf8*^ΔTB^) through two successive rounds of CRISPR/Cas9-mediated targeting in zygotes (Fig. 6C). In the resulting *Irf8*^ΔTB^ mice compared to controls (*Irf8*^WT^), the DC-enriched fraction of the spleen and BM showed a ∼50% reduction in *Irf8* expression (Fig. 6D). Accordingly, the levels of IRF8 protein in *Irf8*^ΔTB^ cDC1s and pDCs were similar to those in heterozygous *Irf8^+/-^* mice (Fig. 6E). IRF8 expression in Ly6C^+^ monocytes and B cells, both of which express intermediate levels of IRF8, were similarly reduced in *Irf8*^ΔTB^ mice (Fig. S7A). IRF8 is strictly required for cDC1 development but is dispensable for pDC development *in vivo* due to compensation by IRF4 (*33*). Despite the observed reduction of IRF8 expression, cDC1 and pDC numbers were unaffected in *Irf8*^ΔTB^ mice (Fig. S7B,C). We therefore turned to FL-BMDC cultures, which force DC development without potential *in vivo* compensatory mechanisms. Similar to primary DCs, cDC1s and pDCs from *Irf8*^ΔTB^ FL-BMDC cultures demonstrated an ∼50% reduction in IRF8 expression (Fig. 6F). In this setting, however, *Irf8*^ΔTB^ cDC1 development was specifically and severely crippled (Fig. 6G). Thus, our results reveal a functional role for *Irf8*-flanking TAD boundaries in enforcing optimal IRF8 expression in DCs and the subsequent development of IRF8-dependent cDC1s.

**Figure 6.**
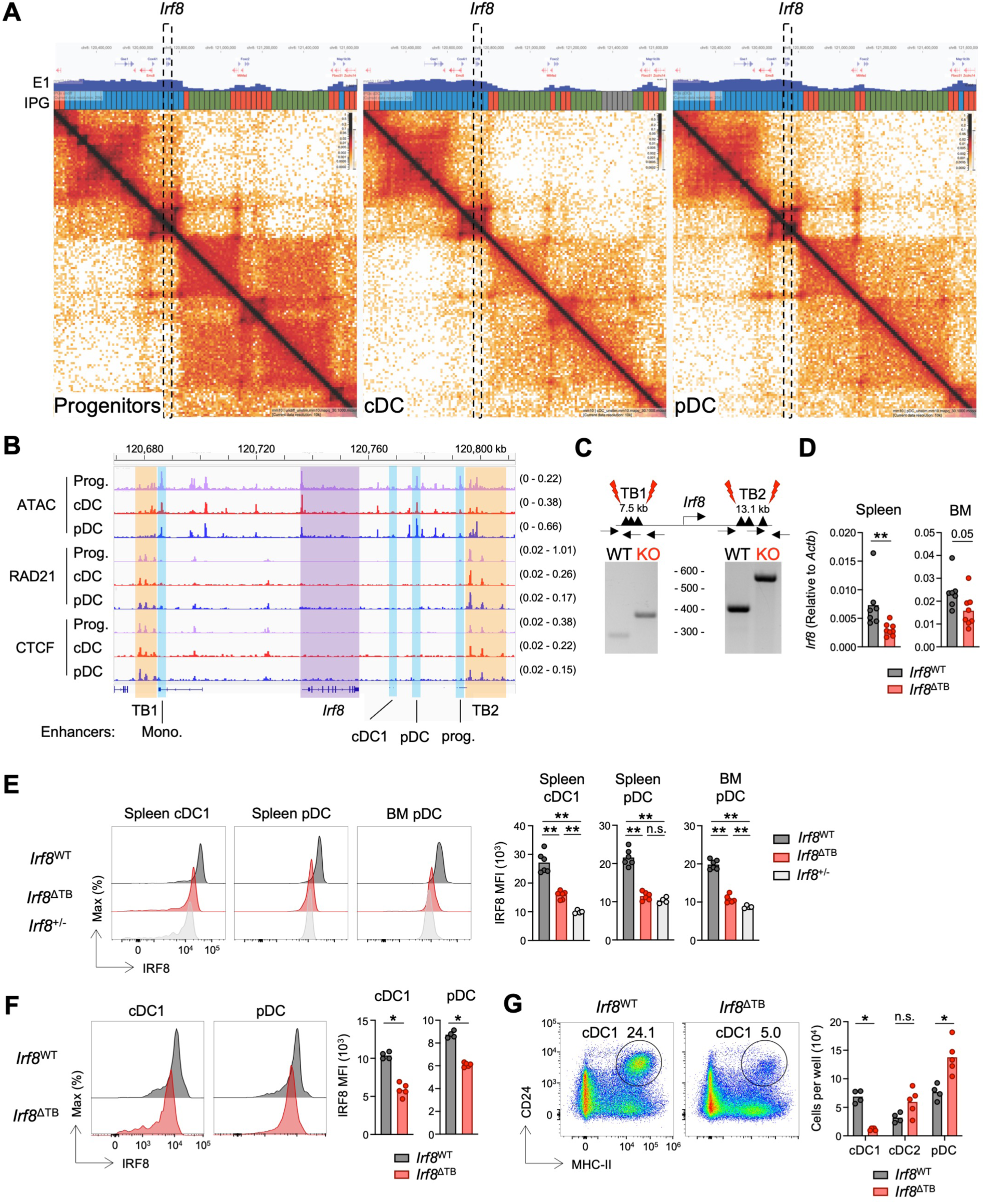
Local Architectural Elements Optimize IRF8 Expression in Developing DCs. (A) Hi-C maps in the vicinity of the *Irf8* locus (highlighed by dashed lines) in Hoxb8-FL progenitors and differentiated cDCs and pDCs. First projected spectral component (E1) and uni-IPG tracks are shown above the Hi-C maps. (B) The chromatin profile (ATAC-seq) as well as Rad21 and CTCF binding profiles (ChIP-seq) in HoxB8-FL progenitors and differentiated cDCs and pDCs. The *Irf8* gene is shaded purple. Cell-type specific enhancers reported to regulate *Irf8* expression in various myeloid populations and their progenitors are highlighted in blue. TAD boundary regions (TB1 and TB2) composed of clusters of CTCF/Rad21 binding sites are highlighted in orange. The scale of each track is indicated on its right. (C) Schematic of the strategy used to excise TB1 and TB2 through sequential rounds of targeting in zygotes to generate *Irf8*^ΔΤΒ^ mice (top). Red lightning bolts indicate sgRNA target sites. Successful deletion of the TB1 and TB2 regions was screened by PCR (bottom) with primer sets indicated by arrows and confirmed by Sanger sequencing. (D) RT-qPCR for *Irf8* (expressed as a ratio relative to that of the housekeeping gene *Actb*) in the DC-enriched fractions of the spleen and BM. Pooled from two independent experiments. (E) Representative histograms (left) and quantification of IRF8 MFI (right) in cDC1s (spleen) and pDCs (spleen and BM) from mice of indicated genotypes. *Irf8^+/-^* mice were used as controls with 50% levels of IRF8. (F) Representative histograms (left) and quantification of IRF8 MFI (right) in cDC1s and pDCs from FL-BMDC cultures derived from the mice of indicated genotypes. (G) Representative flow plots (left) and numbers (right) of DC subsets from FL-BMDC cultures. Symbols represent individual mice, and bars represent mean. Statistical differences were evaluated using Mann-Whitney test. *p < 0.05, **p < 0.01; n.s. not significant.

### CTCF sites control basal and inducible IL-12 production in cDC1s

Finally, we directly tested the role of cohesin-mediated control of gene expression in DCs *in vivo* focusing on its identified key target, IL-12 (Fig. 2A-E). *Il12b* induction in activated macrophages depends on the precise coupling of transcription at the *Il12b* promoter and its HSS1 enhancer located ∼10 Kb upstream (*63, 64*). We thus postulated that the spatial positioning of the *Il12b* locus and its *cis*-regulatory elements would be critical for their coordination. Hi-C analysis revealed that the *Il12b* locus is located within a “private” ∼123 Kb sub-TAD that transitions from B to A genomic compartments during the differentiation of HoxB8-FL progenitors to mature DCs (Fig. 7A), suggesting that developmental changes in chromatin structure prime the *Il12b* locus for subsequent activity. The *Il12b* sub-TAD contains the characterized HSS1 enhancer, and single CTCF/cohesin binding sites constitute its upstream and downstream boundaries (Fig. 7B). To test the role of these sites in *Il12b* regulation, we concomitantly targeted the CTCF binding motifs at each TAD boundary with a single sgRNA in zygotes (Fig. 7C). By PCR and Sanger sequencing, we confirmed 11 bp deletions interrupting the majority of the CTCF motifs at both TAD boundaries (Fig. 7C). The resulting *Il12b*^ΔCTCF^ mice manifested lower levels of IL-12p40 in their serum in the steady state (Fig. 7D). Profilin-induced production of IL-12p40 was similarly impaired; of note, a substantial fraction of *Il12b*^ΔCTCF^ mice showed very little to no IL-12p40 response (Fig. 7D), suggesting that CTCF-binding sites and associated cohesin activity increase the probability (rather than the magnitude) of the activation-induced IL-12 response *in vivo*. *Il12b*^ΔCTCF^ cDC1s enriched from spleens or from FL-BMDC cultures displayed a similar impairment in the steady-state and profilin-induced *Il12b* expression (Fig. 7E,F). This impairment was specific to *Il12b* compared to other inducible cytokines (e.g. *Il6*, *Tnf*), suggesting that the development and activation programs of *Il12b*^ΔCTCF^ cDC1s are otherwise normal (Fig. S7D,E). Collectively, these data reveal a role for CTCF/cohesin sites in the hallmark function of cDC1s, namely IL-12 production.

**Figure 7.**
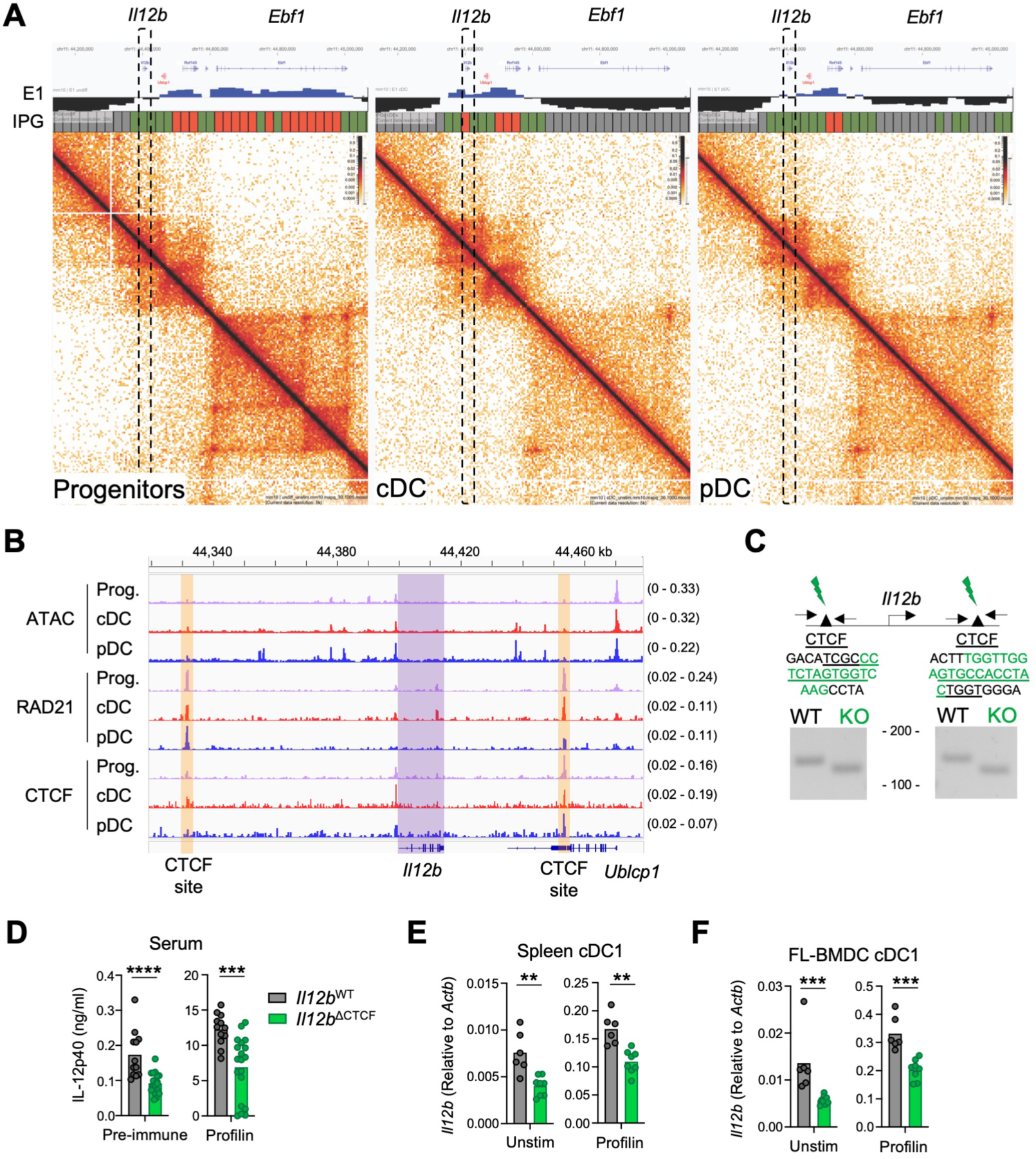
*Il12b* CTCF Sites Control Basal and Inducible *Il12b* Expression. (A) Hi-C maps in the vicinity of the *Il12b* locus (highlighed by dashed lines) in Hoxb8-FL progenitors and differentiated cDCs and pDCs. First projected spectral component (E1) and uni-IPG tracks are shown above the Hi-C maps. (B) The chromatin profile (ATAC-seq) as well as Rad21 and CTCF binding profiles (ChIP-seq) in HoxB8-FL progenitors and differentiated cDCs and pDCs. The *Il12b* gene is shaded purple. The TAD boundaries composed of a single CTCF/Rad21 binding site on either side of the locus are highlighted in orange. The scale of each track is indicated on its right. (C) Schematic of the strategy used to target the CTCF binding sites upstream and downstream of *Il12b* to generate *Il12b*^ΔCTCF^ mice (top). Green lightning bolts indicate sgRNA target sites. Significant deletion of the CTCF binding motif was screened by PCR (bottom) with primer sets indicated by arrows and confirmed by Sanger sequencing (middle). The sequence of the wild-type CTCF binding site is underlined, and the deletion observed in founders is highlighted in green. (D) Quantification of IL-12p40 concentration in serum of indicated mice prior to (left) and 6 hours post challenge with profilin (right). Pooled from two independent experiments. (E-F) RT-qPCR for *Il12b* (expressed as a ratio relative to that of the housekeeping gene *Actb*) in spleen enriched for cDC1 (panel E) and cDC1s purified from FL-BMDC cultures (panel F) of indicated mice that were either unstimulated (left) or stimulated with profilin (right) for 4 hours *in vitro*. Symbols represent individual mice, and bars represent mean. Statistical differences were evaluated using Mann-Whitney test. **p < 0.01, ***p < 0.001, ****p < 0.0001.

## Discussion

Studies over the last decade have described the architecture of mammalian genomes at increasingly higher resolution (*65*) as well as identified the key role of cohesin in its organization (*66*). Furthermore, many groups have characterized the dynamic reorganization of this architecture during embryonic development (*67*) and cell differentiation (*68, 69*). On the other hand, the causal role of cohesin-mediated genome folding in gene expression and the resulting tissue physiology is less well understood. Deletion of the cohesin subunit Nipbl in hepatocytes markedly ablated TADs yet resulted in only modest changes in transcription, with the effect on cell differentiation unknown (*11*). Furthermore, inducible deletion of the cohesin subunit Rad21 in mature macrophages only minimally affected the maintenance of the macrophage transcriptome without activation (*54*). Thymocyte differentiation was shown to be cohesin-dependent entirely due to defective *Tcra* rearrangement, as proper differentiation could be restored by providing thymocytes with a pre-arranged TCR (*70*). On the other hand, major effects of cohesin loss in the developing brain (*71*) or in B lymphocyte responses (*72*) may in part reflect the role of cohesin in cell division.

Here we established both *in vivo* and *in vitro* systems to interrogate the role of cohesin in the differentiation and function of DCs independently of its effect on proliferation. We found that even an incomplete (<50%) deletion of Smc3 from DCs impaired all cohesin-dependent features including cohesin stopping at CTCF sites, CTCF-facilitated interactions (stripes) and cohesin loading at enhancers (fountains). The results revealed a critical role for cohesin in the terminal maturation program of cDCs, particularly of cDC1s. The preferential sensitivity of cDCs vs pDCs to cohesin levels may be attributable to their different life histories: whereas pDCs develop in the BM and emerge as mature cells, cDC differentiation continues in the periphery, driven by tissue-derived signals. Furthermore, the pDC lineage-defining TF *Tcf4* is already expressed in stem cells and multipotent progenitors (*73*) whereas TFs driving cDC differentiation must be induced upon commitment. The observed preferential cohesin dependence of cDC1s vs cDC2s may reflect the fact that cDC1s are a distinct lineage with unique progenitors, driven by multiple specific TFs (i.e. IRF8, NFIL3, ID2, BATF3) (*74, 75*); in contrast cDC2s are thought to be more plastic, a convergent state of differentiation from varied progenitors (*76–79*). Critically, cohesin was required for hallmark cDC1 functions including cross-presentation and antitumor responses in vivo, highlighting its essential functions in immune responses.

Given that the function of DCs is triggered by pathogen-induced activation, we used them to study the role of cohesin in activation-inducible gene expression. Whereas inducible gene expression in macrophages was found to be partially impaired by cohesin loss (*54*), the activation program of cohesin-deficient cDCs was predominantly intact, except for a specific subset of inducible genes. This may be in part explained by our finding that inducible genes are already repositioned into active compartments (IPG) during the DC differentiation process. Furthermore, during DC activation, which proceeds rapidly over just a few hours and without cell division, genome structure is only minimally reorganized (*58, 80*). Consistent with this anticipatory effect, we observed that steady-state cohesin-deficient cDCs exhibited a dysregulated maturation (Mat^ON^) signature, which is comprised of many genes that are further induced during cDC activation. Thus, when establishing cohesin dependence, the process of “priming” via differentiation-dependent chromatin reorganization should be clearly distinguished from true inducibility. On the other hand, we found that the small subset of highly cohesin-dependent inducible genes tends to be farther from active enhancers and are enriched in Polycomb-dependent repressive chromatin marks. By inference, most of the genes of the induction program that are cohesin-independent, have already acquired histone marks of activation after differentiation, and/or are driven by proximal rather than distal enhancers. This is consistent with recent studies in synthetic systems, which showed that cohesin is crucial for establishing distal enhancer-promoter interactions (*5, 6*). Collectively, these data demonstrate that cohesin-mediated genome organization facilitates inducible responses both by “priming” inducible loci during differentiation for their subsequent induction, and by facilitating enhancer-promoter interactions and de-repression during activation.

Lineage-specifying TFs were shown to contribute to the control of genome organization by modulating chromatin looping in lymphocytes (*19, 20, 22*). Myeloid progenitors deficient in IRF8 were shown to have abnormal Hi-C profiles (*58*), whereas subsequent differentiation stages could not be examined. Our genetic analysis of IRF8 revealed its critical contribution to cohesin-mediated genome organization, which could be direct (via cooperation with cohesin or its regulators on the chromatin) and/or indirect (via the enforcement of terminal differentiation). In addition, we undertook an integrative analysis of genome compartments during DC differentiation by generalizing the IPG approach (*55*) for developmental trajectories of Hi-C. This approach revealed that IRF8 was required for transitions between compartments that underlie gene expression changes. Thus, both cohesin and specific TF have essential roles in differentiation: while cohesin is responsible for local chromosome folding, TF drive differentiation and associated chromosome reorganization at all levels, including compartments. Critically, we also uncovered an unexpected reciprocal interaction between the two mechanisms, i.e. the optimization of IRF8 expression by cohesin, via the TAD boundary CTCF elements around the *Irf8* gene itself. The observed strong insulation of the gene encoding an essential TF is likely to represent a common mechanism that ensures robustness of TF-driven cell differentiation programs.

Cohesin-mediated loop extrusion is believed to ensure robust and precise gene expression by facilitating enhancer-promoter interactions within CTCF-demarcated TADs, and by insulating at CTCF boundaries against the spread of enhancer activity to and from adjacent domains (*81*). These conclusions have been drawn primarily from studies of CTCF sites within well-characterized networks of *cis*-regulatory elements in various developmental contexts (*82–85*), including lymphocytes (*86, 87*). Here, we studied the in vivo role of CTCF binding sites in loci that are critical for cDC1 development (*Irf8*) and function (*Il12b*). In the case of *Irf8*, the removal of its flanking TAD boundaries reduced *Irf8* transcription and IRF8 protein expression ∼2-fold, limiting cDC1 differentiation *in vitro*. Mutagenesis of *Il12b*-flanking CTCF sites decreased both basal and inducible *Il12b* expression ∼2-fold, leading to a defective and heterogenous *in vivo* cDC1-driven IL-12 response to profilin. These findings beget several conclusions about the function of CTCF/cohesin. First, rather than acting as binary on-off switches, CTCF sites promote optimal gene expression. This is in contrast to enhancer deletions, which in certain scenarios (e.g. *Irf8* +32 Kb enhancer) can completely ablate lineage development (*60*). Thus, even in the absence of their nearest CTCF site, enhancers may still be capable of reaching their target promoters, perhaps via cohesin-mediated extrusion not targeted by CTCF. Second, in addition to optimizing the magnitude of gene expression, CTCF/cohesin-mediated regulation also controls its robustness, particularly under inducible conditions. This was observed most clearly in the case of *Il12b* CTCF site mutagenesis, in which the inducible *in vivo* IL-12 response was not only diminished but also rendered more variable. A recent report identified the coupling of enhancer and promoter bursting as the key cohesin-dependent parameter during inducible responses in macrophages (*63*). Given that coupling is an intrinsically probabilistic event at the level of individual alleles, this may create a thresholding effect wherein a certain amount of coupling is required for a productive inducible response.

In conclusion, our studies demonstrate the essential role of cohesin-mediated chromatin organization in cell differentiation, inducible gene expression and ultimately in efficient immune responses in vivo. They also reveal both distinct and cooperative pathways of chromatin regulation by cohesin and transcriptional master regulators such as IRF8, as well as an unexpected role of the former in the control of the latter. Further studies in other cell types within and beyond the immune system should help elucidate the full spectrum of cohesin-mediated control of organismal functions.

## Acknowledgments

We thank Maxim Imakaev for the initial analysis of Hi-C data, and Thomas Reimonn for helpful discussions. Supported by the NIH grants AI072571, AI154864 and AI164728 (B.R.), GM114190 and HG011536 (L.A.M.), the Damon Runyon Cancer Research Foundation postdoctoral fellowship DRG 2408-20 (N.M.A.), and the Dr. Bernard Levine postdoctoral fellowship program in immunology (I.T., N.M.A.). L.A.M is a Simon Investigator. We acknowledge the use of resources provided by NYU Rodent Genetic Engineering Laboratory (RGEL), Genome Technology Center (GTC), Cytometry and Cell Sorting Laboratory (CCSL), Applied Bioinformatics Facility Laboratories (ABL) and the High Performance Computing Facility (HPCF). RGEL, GTC and ABL are Shared Resources partially supported by NIH grant CA016087 at the Laura and Isaac Perlmutter Cancer Center.

## Author contributions

N.M.A, I.T., J.R. and G.S.Y. performed experiments and analyzed results. A.G., E.E., C.M.L, N.A., A.S. and A.K.-J. developed analytical methods and analyzed results. G.S.Y. provided essential reagents. N.M.A., A.G., L.A.M., and B.R. conceived the project, analyzed results and wrote the manuscript with input from all authors.

## Materials and Methods

### Mice

All mice used in this study were housed and bred under specific pathogen-free conditions at New York University Grossman School of Medicine (NYUGSoM). All animal studies were performed according to the investigator’s protocol approved by the Institutional Animal Care and Use Committee of NYUGSoM. Mice with conditional deletion of *Smc3* were generated by crossing the loxP-flanked conditional allele of *Smc3* (*37*) with DC-specific *Itgax-*Cre (*38*) (*Smc3*^ΔDC^) or with tamoxifen-inducible *Rosa26^CreER/^*^+^ (*88*). Mice with deletion of *Irf8*-flanking TAD boundaries (*Irf8*^ΔTB^) or disruption of *Il12b*-flanking CTCF binding sites (*Il12b*^ΔCTCF^) were generated on C57BL/6N background and back-crossed at least once to C57BL/6 prior to inter-crossing. Unless otherwise specified, experiments were performed with adult mice 6-10 weeks of age. No differences were observed between male and female mice in any experimental system, so mice of both sexes were used throughout this study. Within an individual experiment however, age- and sex-matched mice were used whenever possible.

### Cell Lines

The Flt3L-secreting clone of the B16 melanoma cell line (B16-Flt3L) (*89*) was cultured in DMEM DMEM supplemented with 10% FCS, 1% L-glutamine, 1% sodium pyruvate, 1% MEM-NEAA, 1% penicillin-streptomycin and 55 μM 2-mercaptoethanol. B16-Flt3L cells were expanded and grown to confluency, and supernatants were collected for Flt3L used in the differentiation of HoxB8-FL and FL-BMDC cultures. The OP9 cell line transduced with a retrovirus encoding GFP and Notch ligand DL1 (OP9-DL1) (*90*) was cultured in MEM-α supplemented with 20% FCS and 1% penicillin-streptomycin (OP9 medium). As previously described (*57*), the murine progenitor HoxB8-FL cell line (*56*) was maintained in the progenitor state by culturing in RPMI supplemented with 10% FCS, 10% B16-Flt3L supernatant, 1% L-glutamine, 1% penicillin-streptomycin, and 10 μM β-estradiol (Sigma-Aldrich). To induce progenitor differentiation into mature DCs, the cells were washed twice and replated in RPMI supplemented with 10% charcoal-stripped serum, 10% B16-Flt3L supernatant, 1% L-glutamine, and 1% penicillin-streptomycin in the absence of estrogen. MC38 colon adenocarcinoma cellswere cultured in RPMI supplemented with 10% FCS, 1% L-glutamine, and 1% penicillin-streptomycin. All cell lines were grown at 37°C in 5% CO_2_. All cells were verified to be *Mycoplasma*-free using the Venor GeM Mycoplasma PCR-Based Detection Kit (Sigma-Aldrich).

### Mouse Generation by CRISPR/Cas9-mediated targeting

sgRNAs that flank *Irf8* TAD boundaries or disrupt *Il12b* CTCF binding sites were identified using Wellcome Sanger Institute Genome Editing (WGE) (https://wge.stemcell.sanger.ac.uk/), CHOPCHOP (http://chopchop.cbu.uib.no/) and Benchling (https://www.benchling.com/). sgRNA efficacy was confirmed prior to use by electroporation into HoxB8-FL cells, followed by the T7 endonuclease I (T7eI) mismatch cleavage assay (IDT Alt-R Genome Editing Detection Kit). Day 0.5 single-cell embryos from C57BL/6N mice were isolated and underwent microinjection with sgRNAs and Cas9 protein at the Rodent Genetic Engineering Laboratory at NYUGSoM. Injected embryos were subsequently transplanted into the oviducts of pseudopregnant recipient mice. The resulting pups were screened by PCR and confirmed by Sanger sequencing to identify those with successful deletion or mutation of the region of interest. Pups heterozygous for deletion or mutation of the desired region were back-crossed to WT C57BL/6 mice, and heterozygous pups were inter-crossed to generate homozygous mice. The CTCF binding site upstream and downstream of *Il12b* were disrupted concomitantly with a pair of sgRNAs, one targeting each binding site. In contrast, the TAD boundaries flanking *Irf8* were removed sequentially. Single-cell embryos from mice homozygous for the downstream *Irf8* TB2 deletion (*Irf8*^ΔTB2^) were microinjected with sgRNAs flanking the upstream TB1 region.

### Mouse procedures

For EdU labeling, Mice were injected i.p. on three consecutive days with 1 mg EdU (Carbosynth) in 200 ul PBS and harvested for analysis one day after the last injection. For SRBC immunization, mice were injected i.p. with 2×10^8^ sheep RBCs (Innovative Research) in 500 ul PBS and harvested for analysis eight days post-immunization. For in vivo cross-presentation, spleen and skin-draining lymph nodes were harvested from BALB/c (H-2^d^) mice, mashed, and pooled together. Cells were pulsed for 10 min at 37°C with a hypertonic solution of 10% polyethylene glycol (Sigma-Aldrich), 0.5 M sucrose (Sigma-Alrich), 10 mM HEPES, and 10 mg/ml chicken ovalbumin (OVA) protein (Invivogen) dissolved in RPMI and then immediately flushed with a hypotonic solution of 40% water/60% RPMI for 2 min at 37°C. Following 2 washes with cold PBS, cells were irradiated with 2000 rad using a Cell-Rad X-Ray irradiator (Precision). Recipient (H-2^b^) mice were injected i.v. (retro-orbital route) with 4×10^7^ cells in 200 ul PBS, and the cDC1-dependent, endogenous OVA-specific CD8^+^ T cell response was analyzed in the spleen on day seven post-injection by flow cytometry using OVA/H-2K^b^ tetramer staining. As a control, recipient mice received irradiated BALB/c splenocytes pulsed with a hypertonic solution lacking OVA protein.

For IL-12 responses, mice were injected i.p. with 100 ng profilin (Sigma-Aldrich) in 500 ul PBS. Blood was collected from mice both before and 6 hours after challenge, and serum separated from whole blood. IL-12p40 and IL-12p70 concentrations in serum were measured by ELISA (Thermo Fisher Scientific Ready-SET-Go! Kit) according to the manufacturer protocol. ELISA plates were read using a FlexStation Microplate reader.

For antitumor responses, MC38 cells were washed once and resuspended in endotoxin-free PBS containing 2.5 mM EDTA. Mice were injected s.c. on both dorsal flanks with 5×10^5^ tumor cells in 100 ul volume. To analyze the response to immunotherapy, on day 6 post-tumor inoculation, mice of each genotype were randomized and injected i.p. with a combination immunotherapy regimen of 200 μg anti-PD-1 blocking antibody (clone 29F.1A12, Bio X Cell) and 200 μg anti-4-1BB agonist antibody (clone LOB12.3, Bio X Cell) or rat IgG2a isotype control antibody (clone 2A3, Bio X Cell) in 200 ul PBS every third day until the experiment endpoint. Tumor growth was measured using a caliper by a researcher who was blinded to genotype and treatment group. Tumor volume was estimated using the formula: Tumor volume ∼ Length x Width^2^ x π/6.

### Mouse Tissue Primary Cell Preparation

Spleens were minced and digested in 1 mg/ml collagenase D (Roche) and 20 μg/ml DNaseI (Roche) in RPMI supplemented with 10% FCS for 30 min at 37°C. Digested splenic tissue was subjected to red blood cell (RBC) lysis (BioLegend). Cleaned femur and tibia bones were crushed with mortar and pestle. The released bone marrow was subjected to RBC lysis. Skin-draining lymph nodes (bilateral inguinal and brachial nodes) were collected and lymph node capsules pierced with sharp forceps. Lymph nodes were digested in 1 mg/ml collagenase D in HBSS with calcium and magnesium for 30 min at 37°C. Lungs were minced and digested in 1 mg/ml collagenase IV (Worthington Biochem) in RPMI supplemented with 10% FCS for 30 min at 37°C. Digested lung tissue was subjected to RBC lysis. Ears were collected and separated into dorsal and ventral halves. Each half was placed with the dermis layer down in digestion media (RPMI supplemented with 250 μg/ml Liberase TL (Roche) and 125 μg/ml DNase I) for 90 min at 37°C. Halfway through the digestion, the ear skin was minced with scissors.

### Flow Cytometry and Cell Sorting

Single-cell suspensions were blocked with TruStain FcX anti-mouse CD16/CD32 (BioLegend) and subsequently stained with fluorophore-conjugated or biotinylated antibodies (BioLegend, Thermo Fisher Scientific, BD Biosciences). Where indicated, some or all of the following non-DC markers (TCR-β, CD3, CD19, NK1.1, NKp46, Siglec-F, Ly6G) were stained with antibodies conjugated to the same fluorophore (lineage dump) to exclude non-DCs in flow cytometric analysis of DC populations. Intracellular staining was performed by fixing and permeabilizing with the eBioscience Foxp3 / Transcription Factor Staining Set (Thermo Fisher Scientific). EdU positive cells were stained with Click-iT Plus EdU AlexaFluor 488 Flow Cytometry Assay Kit (Thermo Fisher Scientific) according to the manufacturer protocol. eBioscience Fixable Viability Dye eFluor 506 (Thermo Fisher Scientific) was used to exclude dead cells in some experiments. Samples were acquired on Attune NxT (Thermo Fisher Scientific) using Attune NxT software and data analyzed with FlowJo software v10 (Tree Star). Cell sorting was performed on FACSAria II (BD Biosciences) at the Cytometry & Cell Sorting Laboratory at NYUGSoM. Flow cytometry was performed on sorted samples to confirm purity > 95% for populations of interest.

### Cell Enrichment

To facilitate sorting of primary splenic DCs (e.g. for RNA-seq or ATAC-seq), a single-cell splenocyte suspension was enriched for DCs by negative depletion. Briefly, splenocytes were stained with biotinylated antibodies against TER-119, TCR-β, CD3ε, NK1.1, CD19, and Ly6G (Thermo Fisher Scientific and BioLegend), incubated with streptavidin microbeads (Miltenyi Biotec) and passed through LS columns attached to MidiMACS or QuadroMACS separators (Miltenyi Biotec) according to manufacturer protocol. In experiments in which RNA was extracted from enriched DC subpopulations for RT-qPCR without sorting, a more stringent depletion was used. For enrichment of spleen pDC/cDC1s, additional biotinylated antibodies against CD90.2, F4/80, CD11b, CD138, IgD, IgM, and SIRPα were added to the above staining cocktail. For enrichment of bone marrow pDCs, enrichment was performed using the more stringent cocktail above on LD columns. For enrichment of cDC1s alone, B220 was also added to the stringent cocktail of biotinylated antibodies.

### FL-BMDC cultures

Primary mouse bone marrow cells were plated at a density of 2×10^6^ cells per well in 2 ml of DMEM supplemented with 10% FCS, 10% B16-Flt3L supernatant, 1% L-glutamine, 1% sodium pyruvate, 1% MEM-NEAA, 1% penicillin-streptomycin, and 55 μM 2-mercaptoethanol (DC medium) in 24-well plates. Cells were cultured at 37°C for 8 days without replating and collected by scraping the bottom of the well. To induce recombination of the floxed *Smc3* allele in *Rosa26^CreER/^*^+^ *Smc3^fl/fl^* cells, 4-hydroxytamoxifen (4-OHT) (Sigma-Aldrich) was spiked into cultures at day 3 of differentiation to a final concentration of 500 nM.

### Notch-driven FL-BMDC cultures

Primary mouse bone marrow cells were initiated as above. On day 2 of differentiation, OP9-DL1 cells were treated with 10 μg/ml mitomycin C (Sigma-Aldrich) for 2 h, harvested, washed three times, and 9×10^4^ cells plated in 1 ml OP9 medium in 24-well plates. On day 3 of differentiation, differentiating bone marrow cells were harvested and 1 ml added per well of OP9-DL1 stromal cells, from which OP9 media was aspirated just prior. On the same day, 4-OHT was spiked into the cultures to a final concentration of 500 nM to induce recombination of the floxed *Smc3* allele in *Rosa26^CreER/^*^+^ *Smc3^fl/fl^* cells. Co-cultures were incubated for 5 more days at 37°C and collected by scraping the bottom of the well. Harvested cultures were passed through 70 μm filters to exclude most OP9-DL1 stromal cells during analysis.

### In Vitro Stimulation

For assays testing *Smc3*^Δ^ cDC1 function by ELISA, FL-BMDC cultures (2×10^5^ cells per well) were stimulated with either 100 ng/ml profilin or 1 μM CpG-B/ODN1668 (Invivogen), or left unstimulated as a control in 200 ul DC medium in 96-well plates. Supernatant was collected 24 hours later for analysis of IL-12p40 and IL-6 production by ELISA (Thermo Fisher Scientific Ready-SET-Go! Kit) according to the manufacturer protocol. ELISA plates were read using a FlexStation Microplate reader.

For assays characterizing *Smc3*^Δ^ cDC activation by RNA-seq, FL-BMDC cultures were stimulated as above with a combination of 1 μM CpG-A/ODN2216 and 1 μM CpG-B or left unstimulated as a control. Cells were collected 6 hours later and sort purified for cDC1 and cDC2 subsets, followed by RNA extraction. Successful cDC activation was confirmed by flow cytometric analysis of CD69 upregulation.

For assays testing *Il12b*^ΔCTCF^ DC IL-12 production by RT-qPCR, splenic DCs (enriched for cDC1) or FL-BMDC cultures (sort purified for cDC1) were stimulated with either 100 ng/ml profilin or left unstimulated as a control for 4 hours, followed by RNA extraction.

### RT-qPCR

RNA was extracted from MACS-enriched or sort purified cells using the RNeasy Mini Kit and the accompanying RNase-Free DNase Set (Qiagen) according to the manufacturer protocol. cDNA was prepared by reverse transcription of extracted RNA using the iScript cDNA Synthesis Kit (Bio-Rad Laboratories), and qPCR of cDNA was performed using the KAPA SYBR FAST qPCR Kit (Roche) according to the manufacturer protocols. Transcript levels of the indicated genes of interest are expressed as a ratio relative to the housekeeping gene *Actb* (encoding β-actin) through the formula 2^-(C_t_(*Gene*) - C_t_(*Actb*))^. qPCR was run on a QuantStudio 3 Real-Time PCR System (Thermo Fisher Scientific).

### DNA Extraction

DNA was extracted from sgRNA-transfected HoxB8-FL cells (for testing of sgRNA efficiency by T7eI assay) or from sort-purified CD11c^+/-^ splenocytes from *Smc3*^ΔDC^ mice (for confirming recombination of floxed *Smc3* allele by PCR) using the NucleoSpin Tissue kit (Takara Bio) according to the manufacturer protocol.

### Immunofluorescence

FL-BMDCs were loaded into Epredia Cytofunnels (Thermo Fisher Scientific) and concentrated onto Epredia Cytoslide microscope slides (Thermo Fisher Scientific) using a cytocentrifuge (Thermo Fisher Scientific). Slides were fixed with 4% paraformaldehyde (Electron Microscopy Sciences) in PBS for 10 min and blocked with PBS + 5% BSA (Thermo Fisher Scientific) + 0.3% Triton X-100 (Thermo Fisher Scientific) for 30 min. Slides were then stained with rabbit anti-SMC3 (clone D47B5, Cell Signaling Technology), AlexaFluor 488 goat anti-rabbit IgG (H+L) (Thermo Fisher Scientific), and DAPI. Images were captured using an EVOS microscope. For better visualization, the color intensity of images was enhanced slightly but consistently on whole images of all slides using Microsoft Powerpoint.

### Western Blot

FL-BMDC cell pellets at day 8 post-differentiation were lysed on ice for 30 min in RIPA Lysis and Extraction Buffer (Thermo Fisher Scientific) supplemented with Halt Protease and Phosphatase Inhibitors (Thermo Fisher Scientific). Samples were boiled for 10 min in the presence of SDS sample loading buffer (BioRad). Lysates of equal protein quantities, as determined by the Pierce BCA Protein Assay Kit (Thermo Fisher Scientific), were analyzed by SDS-PAGE followed by Western blotting with rabbit anti-SMC3 or mouse anti-β actin as a loading control. As a reference point for Smc3 protein levels, protein extracted from the FL-BMDC cultures of *CD11c*-Cre *Smc3^fl/^*^-^ mice were used.

### Generation of Irf8^Δ^ HoxB8-FL Cells

Cas9 ribonucleoproteins (RNPs) were prepared by incubating 250 pmol IRF8 sgRNA, 250 pmol CD45 sgRNA, 75 pmol *S. pyogenes* Cas9 Nuclease V3, and 100 pmol Alt-R Cas9 electroporation enhancer (all from IDT) together for 15 min at room-temperature. Control Cas9-RNPs were prepared containing 250 pmol non-targeting control crRNA and 250 pmol tracrRNA instead of 250 pmol IRF8 sgRNA. HoxB8-FL cells (1×10^6^) were resuspended in 100 ul Opti-MEM I reduced serum medium (Thermo Fisher Scientific), to which Cas9-RNPs were added. Cells were electroporated with 4D-Nucleofector X Unit (Lonza) using program CM-137. Transfected HoxB8-FL cells were passaged every other day until day 6 post transfection, at which time HoxB8-FL progenitors were differentiated into mature DCs. Successful transfection was confirmed by loss of CD45 expression on both IRF8-deficient (*Irf8*^Δ^) and control HoxB8-FL cell lines.

### RNA Sequencing

RNA was extracted from up to 5×10^4^ sort-purified cells using the RNeasy Mini Kit and the accompanying RNase-Free DNase Set according to the manufacturer protocol. RNA quality and quantity was assessed by Agilent BioAnalyzer. cDNA libraries were prepared using the low input Clontech SMART-Seq HT with Nxt HT kit according to the manufacturer protocol, and paired-end 50 bp sequencing was performed on Illumina NovaSeq 6000 at the Genome Technology Center at NYUGSoM.

### ATAC Sequencing

Up to 5×10^4^ sort-purified cells were washed and prepared as previously described (*91*). Paired-end 50 bp sequencing was performed on Illumina NovaSeq 6000 or Illumina HiSeq 4000 at the Genome Technology Center at NYUGSoM.

### Hi-C

Sort-purified cells (∼1-5×10^6^) were cross-linked for 10 min in 1 ml of 2% methanol-free formaldehyde (Thermo Fisher Scientific) in PBS + 3% BSA. Crosslinking was reversed by addition of glycine (Thermo Fisher Scientific) to 0.125 M for 5 min, followed by incubation on ice for 15 min. Libraries were prepared using the Arima Hi-C kit according to manufacturer protocol and paired-end 100 bp sequencing of libraries was performed on Illumina NovaSeq 6000.

### ChIP-seq

Sort-purified cells (∼3.5-4.5×10^6^) were cross-linked with 1% methanol-free formaldehyde in PBS for 8 min followed by quenching with 0.125 M glycine for 5 min. Cells were lysed and nuclei isolated using the Covaris TruCHIP Kit, and chromatin was sheared using a Covaris M220 ultrasonicator. ChIP was performed using 4 μg of rabbit anti-CTCF polyclonal antibody (Millipore) or rabbit anti-mouse/human Rad21 polyclonal antibody (Abcam) and Dynabeads Protein A (Thermo Fisher Scientific). Libraries were prepared using the KAPA HyperPrep Kit (Roche). As a control, libraries were also prepared from non-immunoprecipitated “input” DNA from each cell type. Paired-end 50 bp sequencing was performed on Illumina NovaSeq 6000 at the Genome Technology Center at NYUGSoM.

### CUT&RUN

CUT&RUN was performed on ∼3×10^5^ sort-purified cells using the CUTANA ChIC/CUT&RUN Kit (Epicypher) according to manufacturer protocol. Briefly, cells were permeabilized with buffer containing 0.05% digitonin. After cells were bound to activated Concanavalin A (ConA)-coated beads, cells were bound with 4 μg of the following antibodies (or 0.5 μg for H3K4me3), which was used to guide cleavage by pAG-MNase: anti-histone H3K4me3 mixed monoclonal (Epicypher), anti-histone H3K4me1 polyclonal (Abcam), anti-histone H3K9me3 polyclonal (Abcam), anti-histone H3K27Ac polyclonal (Abcam), anti-histone H3K27me3 polyclonal (Millipore) and rabbit IgG isotype control polyclonal (Abcam). Libraries were prepared with NEBNext Ultra II Library Prep Kit. Paired-end 50 bp sequencing was performed on Illumina NovaSeq 6000 at the Genome Technology Center at NYUGSoM.

### Statistical analysis

For graphs, data are shown as mean overlaid with values from individual replicates. Statistical differences were evaluated using a non-parametric Student’s t test with Mann-Whitney analysis, unless otherwise indicated. Statistical differences in mice with < 250 mm^3^ tumor volume were determined with the log-rank (Mantel-Cox) test. P values less than or equal to 0.05 were considered significant, and significance was assigned according to the following breakdown: *p < 0.05, **p < 0.01, ***p < 0.001, and ****p < 0.0001. GraphPad Prism software v8 and v10 were used for all graphing and statistical calculations except for gene expression and chromatin analysis, for which R was used. The number of experiment repetitions are indicated in the respective figure legends.

### Computational analysis of sequencing datasets

#### ChIP-Seq data analysis

Data was processed with nf-core (*92*) chipseq nextflow pipeline version 2.0.0 [https://nf-co.re/chipseq/2.0.0/] with parameters --paired_end -- narrow_peak. Briefly, mapping was done by bwa mem v0.7.17-r1188 to the reference mm10 genome, and the peak calling was done by MACS2 v2.2.7.1 in narrow-peak regime. Inputs for normalization were specified in the samplesheet for the nextflow pipeline according to the software documentation.

#### ATAC-Seq Data Analysis

Mapping ATAC-Seq was done with nf-core (*92*) atacseq version 2.1.2 [DOI:10.5281/zenodo.2634132]. Briefly, sequencing reads were mapped to the reference genome (mm10) using the Bowtie2 (v2.5.1) (*93*) and duplicate reads were removed using Picard tools (v.3.1.1) [broadinstitute.github.io/picard/]. Low quality mapped reads were removed from the analysis. The read per million (RPM) normalized BigWig files were generated using BEDTools (v.2.31.1) (*94*) and the bedGraphToBigWig tool (2.9 bbi v.4). Peaks were called with MACS2 in narrow peak mode (*95*).

#### Calling the Chromatin States with ChromHMM

We started with mapped data for CUT&RUN for H3K27ac, H3K4me1, H3K4me3, H3K27me3, and H3K9me3 in pDC, cDC1, cDC2, and undifferentiated cells. We then calculated coverage in 200 bp bins of the genome with a ‘stackup’ function of pybbi [https://zenodo.org/doi/10.5281/zenodo.10382980], a Python wrapper for Big Binary Indexed file tools (*96*). We then binarized the signal by running ChromHMM v1.23 (*97*) with default parameters and learned a set of ChromHMM models with the number of states ranging from 5 to 10.

We then performed manual annotation of the resulting chromatin states and selected the model with 8 states as the best recapitulating biologically relevant combinations of histone modifications: TSS1 and TSS2 are two active transcription states enriched in H3K4me3, differing by the levels of H3K27ac; enhancer is a state with enriched H3K4me1 and mildly increased H3K27ac; BivTSS state was enriched in both H3K27me3 and H3K4me1 and to a lesser extent H3K4me3 and H3K27ac; repressed state enriched solely in H3K27me3; inactive state enriched in H3K9me3; and two types of quiescent chromatin, where all the histone marks are depleted (Supplemental Methods Fig. 1A).

#### RNA-Seq Analysis

For general differential gene expression analysis, we used edgeR (*98*) to perform pairwise comparisons between the following conditions: 1) unstimulated versus stimulated control cDC1s, 2) unstimulated versus stimulated control cDC2s, 3) unstimulated versus stimulated *Smc3*^Δ^ cDC1s, 4) unstimulated versus stimulated *Smc3*^Δ^ cDC2s, and 5) control versus *Irf8*^Δ^ DCs. For each comparison, all available replicates were considered, and within each condition, convergence of replicates was confirmed by PCA. Genes were filtered with a minimum number of 100 RNA-Seq counts in at least some samples. The data was normalized for coverage, its dispersion estimated and the glmQLFit model (quasi-likelihood negative binomial generalized log-linear model) was fitted to the data. The logarithm to the base 2 of fold change (logFC) was used in the expression and p-value of this model.

To define “Smc3-dependent genes” during cDC activation, we considered genes that meet both the following conditions: 1) significantly change in expression in WT cDCs upon stimulation (log_2_FC > 2, p < 0.01), and 2) fail to change in expression to the same extent in *Smc3*^Δ^ cDCs upon stimulation (WT log_2_FC / *Smc3*^Δ^ log_2_FC > 2). “Smc3-independent genes” were defined as those that satisfy only condition 1. All remaining genes were deemed “non-inducible” and retained as a control. “Smc3-dependent genes” were defined separately for cDC1s and cDC2s.

For each gene, we determined the position of its transcriptional start site (TSS) and the CTCF-insulated neighborhood housing its TSS (Supplemental Methods Fig. 1B). The boundaries of CTCF-insulated neighborhoods were defined as the nearest CTCF ChIP-Seq peaks in cDCs, with CTCF motifs in convergent orientation (left boundary is the closest upstream CTCF on the plus strand, while the right boundary is the downstream CTCF on the minus strand). For each gene, all enhancers within its insulated neighborhood were identified and the enhancer closest to the TSS selected for further analysis.

#### Hi-C Data Mapping

Hi-C data was processed with distiller-nf version 0.3.3 with default parameters. Only pairs with both reads mapped with MAPQ>=30 were considered for the final contact matrices. Briefly, reads were mapped with bwa mem [https://arxiv.org/abs/1303.3997] to mm10 reference genome, parsed into contact pairs by pairtools (*99*), then deduplicated and filtered by MAPQ. The data was then loaded into cooler-formatted contact matrices (*100*), aggregated to multiple resolutions and iteratively corrected (*59*). We always used iteratively corrected data for downstream analysis of Hi-C maps.

Hi-C maps were visualized in online service Resgen [https://resgen.io/], a genome browser based on the HiGlass interactive web-based visualization tool (*101*).

#### Hi-C Analysis

For PCA, pairwise Stratum-adjusted Correlation Coefficients (SCC) (*102*) were constructed between all Hi-C matrices in the analysis at 50 Kb resolution, with the maximum distance for contacts set to 1 Mb to exclude noisy long-range interactions. PCA implemented in scikit-learn was then applied (*103*).

To construct P(s) curves, Open2C library cooltools was used (*104*), in particular its “expected” module. Average interaction frequencies as a function of genomic separation between pixels were first calculated with the function *expected_cis*. As a result, the slopes of the P(s) curves were also calculated. Next, P(s) curves were aggregated per region over exponentially increasing distance bins with the *logbin_expected* function to achieve a better spread of data points over genomic separations. Finally, by-region log-binned expected and slopes were combined into genome-wide averages, handling small chromosomes and “corners” optimally, robust to outliers. For the spread, 90% confidence intervals were calculated for each genomic separation range. Finally, the significance of differences between distributions of slopes at 50-150 Kb was calculated with Moods median test and the Levene test for variance equivalence.

To construct average pileups, the *cooltools* “pileup” module (*104*) was used to obtain 300 Kb square snippets of Hi-C maps around the forward-oriented CTCF peaks. The peak positions were defined by MACS2 peak calling (*105*), and the orientation was defined as positive if all the motifs overlapping the peak were mapped to the plus strand of the reference genome. The motifs were called with JASPAR (*106*) UCSC tracks code wrapper for PWMScan (*107*) as was done previously (*108*), with the MA0139 reference motif and GC content calculated for the reference genome (A 0.291, T 0.291, G 0.208, C 0.208). Threshold p-value was set to 1e^-4^ and r to 0.5.

To assess the level of cohesin activity, we took Rad21 degron Hi-C data (*51*) as the maximum (untreated) and minimum (auxin-treated) possible level of cohesin activity, re-mapped the data with *pairtools* and *distiller-nf* (*99*) and calculated P(s) curves and their slopes (same as for our datasets).

The local compaction by cohesin (or P(s) “shoulder” most prominent in the P(s) derivative plot) is observed as a region of the P(s) curve with a shallow slope at separations of ∼100 Kb both in our and (*51*). Thus, the P(s) slope at 100 Kb was considered as the baseline signature of fully functional and expressed cohesin.

Next, *in silico* mixtures of cohesin-depleted and untreated cells were created by sampling the contacts of (*109*) Hi-C maps and combining them at ratios corresponding to N% effectiveness degradation N=10, 20, …, 90. P(s) and its slope were estimated *in silico* mixtures. The slope at 100 Kb was then calculated and plotted as a function of cohesin degradation effectiveness (N). Indeed, the drop in the slope becomes more pronounced as the contribution of cohesin-depleted cells becomes more prominent in the mixtures. This plot served as a “ruler” for the assessment of the level of cohesin activity, which we used to estimated the drop in the P(s) slope at 100 Kb for our datasets and plotted them against the “ruler” derived from (*51*).

#### Fountain Calling in Hi-C Data with Autoencoder

To automatically mark fountains, a variational autoencoder based on 2d-convolutional blocks (Supplemental Methods Figure 1C-H) was used. The model was trained to reproduce high-quality Micro-C maps (*110*) encoding the 640-Kb Hi-C windows into 128-dimensional vectors. As an input for autoencoder, we used Hi-C observed over expected maps sliced along the main diagonal and rotated 45 degrees. Next, only the first part of the model, called the encoder, was used, compressing the input data into a latent representation. We then used the resulting latent representation to determine the similarity of Hi-C patterns to typical 3D genome patterns. For the fountain pattern, we used a mask of zeros containing a triangular pattern of ones in the center, expanding upward. This mask was fed to the trained Hi-C encoder; the output was a vector of the “hidden representation” of the fountain. We then projected the representation of each genomic loci onto the vector corresponding to the fountain mask. This projection served as an autoencoder-based fountain score, a measure of similarity of Hi-C map of each genomic locus to the fountain pattern. We then detected peaks of fountain score with cooltools peak calling function (*104*), and considered the peaks that had the fountain score falling into top 10% as the potential fountain locations.

#### Feature Strength Assessment

For convolutional fountain score, we first created the convolution kernel with fontanka’s (*53*) double triangle mask with angle of 45°, where all the values are zero except the values in the fountain-like shape in the middle of the map. We then multiplied each snippet of Hi-C map by double triangle kernel and used the sum of the multiplied matrices as a measure of fountain strength for average pileups at potential fountains.

#### Unified Interaction Profile Groups

To detect the regions with similar patterns of long-range interactions, we switched to spectral analysis of Hi-C maps, which was previously used to identify two types of chromatin, or compartments (*59*), and more detailed interaction profile groups (*55*).

Briefly, we aimed to obtain a set of leading spectral components to characterize the cluster structure between 25 Kb-genomic regions. Due to significant data sparsity at this resolution, we refrained from using trans-interactions and analyzed more information-rich and dense cis-interactions of chromosomes.

In our analysis, we aimed to compare the patterns of long-range interactions between different cell types progressing in the lineage of dendritic cell development and activation. However, the spectral decomposition of individual datasets results in independent eigenvectors and, thus, independent cluster structure between individual datasets (*55*).

To test for this artifact, we first performed independent spectral clusters analysis in each dataset.

##### Hi-C map preprocessing

We first iteratively corrected cis-Hi-C maps, truncated the bottom 1% and top 99.95% of the signal in pixels, and replaced the first two diagonals with zeros. We then normalized the maps by expected, removed all bad bins, and scaled the columns and rows of the resulting matrix to 1, as done in (*55*).

##### Independent spectral clustering of cis-Hi-C maps

Eigendecomposition of pre-processed matrices was done without the prior mean-centering (*55, 59*). We then compared each leading spectral components to the GC content and flipped the sign of the spectral component if we observed a negative correlation. This allowed us to compare the embeddings of each Hi-C matrix and observe substantial variability in the resulting structures of the embedded space (Fig. S5B). In particular, the embedding structure for undifferentiated cells diverged the most, with the data looking flipped relative to other datatypes; in pDCs from FL-BMDC cultures, we observed an additional “branch” of active chromatin. Thus, we concluded that comparing cluster structures between cell types may be hindered due to independent decomposition, and we switched to the joint decomposition of Hi-C maps of DC lineage (below, and Fig. 4B).

##### Joint spectral clustering of cis-Hi-C maps

We first detect bad bins in each cis Hi-C map and remove the union of all bad bins from the analysis. We then pre-process each Hi-C matrix as described above to obtain normalized non-zero-centered maps. Next, we adapted an extension of principal component analysis for multiple datasets called STATIS (*111*). Briefly, this method (1) accounts for relationships between different datasets, (2) reveals consensus, a common structure between these datasets, and (3) projects each dataset onto consensus to analyze the datasets jointly. STATIS has two crucial steps:

(1) Generalized Singular Value Decomposition (GSVD) that defines the relationship between datasets and reveals the optimum weights of each dataset. The weights will introduce more importance to the maps representing the group, patterns of long-range interactions of DCs, and less importance to more unique maps (such as progenitors).
(2) Generalized PCA of stacked Hi-C maps with previously defined weights as constraints. This approach will reveal a single consensus representative of all maps of DCs, providing unified coordinates for DC long-range interactions.

Finally, we project each individual dataset onto consensus, compare the resulting embeddings, and perform k-means clustering for the rows containing embeddings for all DC datasets and the chromosomal regions (Fig. 4B). The resulting embeddings are similar to those obtained by independent decomposition but are much more consistent between cell types (Fig. S5B). This procedure was repeated for each autosome of mouse at 25 Kb resolution.

##### Choice of the number of leading components and the number of clusters

To choose the number of components, we analyzed the spectrum of decomposed stacked Hi-C matrices. The number of singular values was usually under 10 for different chromosomes, and we selected five leading components as a representative number. The increase in the number of leading components leads to minor changes in the downstream analysis.

For the number of clusters, we calculated silhouette scores for clusters ranging from 2 to 21 and the number of leading components ranging from 2 to 15. Visual analysis revealed a minor contribution of the number of components and a significant drop in silhouette score at around 5 clusters (on average between different chromosomes).

We further assessed the enrichment of each available histone modification from CUT&RUN and confirmed that five clusters produce a reasonable balance between biological interpretation, clustering metrics, and reproducibility between different chromosomes.

Although decomposition and clustering were performed independently for each chromosome, we ensured that the resulting clusters had similar biological interpretations between chromosomes. Each chromosome had one major euchromatic cluster enriched in H3K27ac, H3K4me3, and H3K4me1, which we labeled A1, and one less enriched euchromatic cluster that we called A2. There were usually two clusters that we name “intermediary”, B1 and B2, with less enrichment of active histone marks and an elevated level of H3K27me3. Finally, there was always a state with major enrichment of H3K9me3 and no other chromatin marks that we called B3, or the most heterochromatic one. We observed only a mild degree of variability of these patterns (such as an increase in H3K27me3 in the active states for some chromosomes), and merged the resulting annotations between all chromosomes for interpretation.

##### Technical implementation

Pre-processing and independent spectral clustering were implemented by a custom Python library based on the Open2C inspectro tool (*55*) [https://github.com/open2c/inspectro]. Joint spectral clustering was done by pySTATIS implementation [https://github.com/mfalkiewicz/pySTATIS] of the STATIS tool (*111*). Clustering was done by sklearn v.1.1.3 KMeans with k-means++ initialization, 100 independent initializations, maximum iterations of 1000 and tolerance of 0.00001. Annotation of each genomic region falling into specific cluster was done by bioframe (*112*). The code is posted and publicly available at github [https://github.com/mirnylab/dendritic_cells].

---

**Figure S1.**
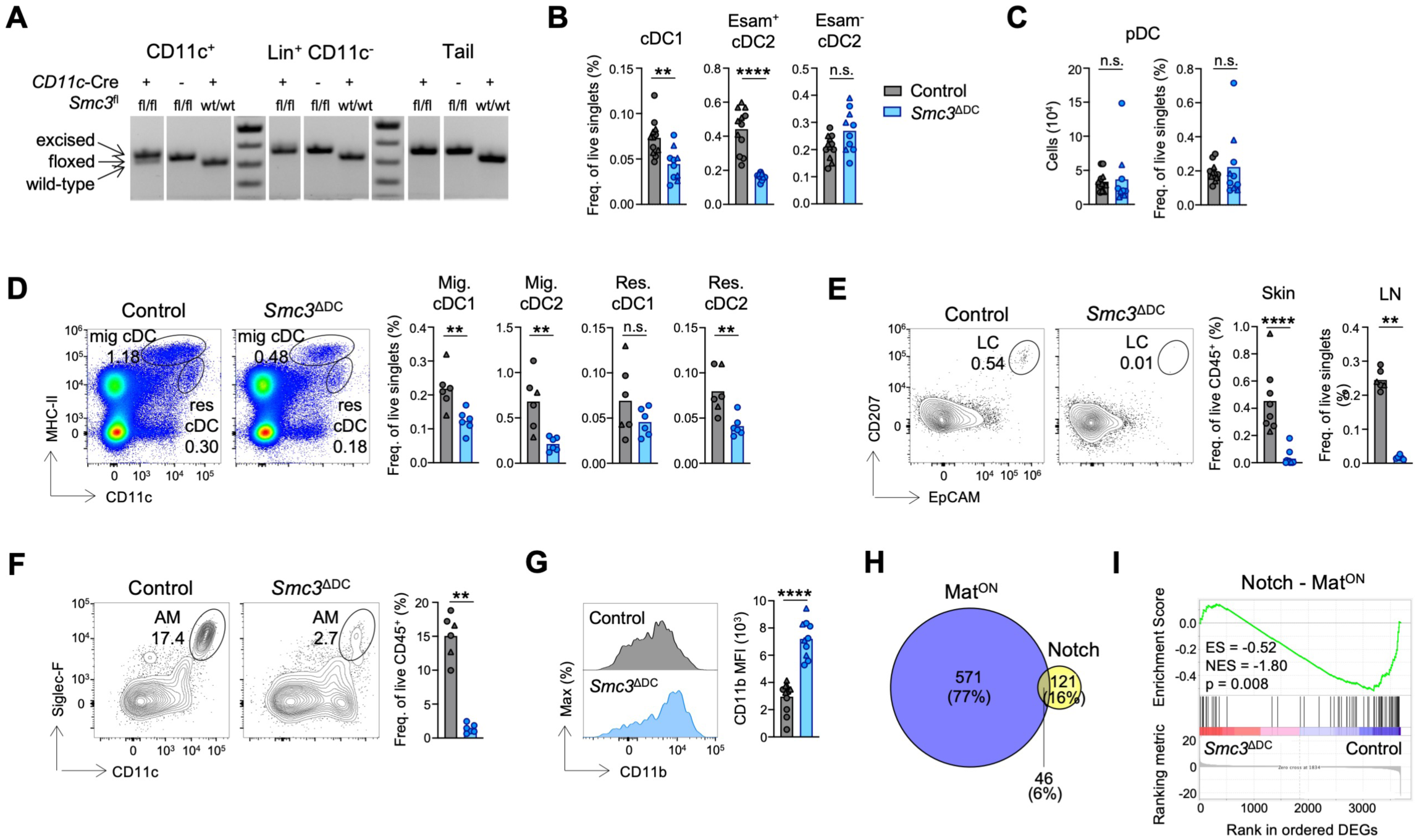
Cohesin is Required for *In Vivo* cDC Differentiation. (A) PCR to detect the recombined *Smc3* floxed allele in purified splenic Lin^-^ CD11c^+^ cells, Lin^+^ CD11c^-^ cells, and tail DNA of indicated mice. (B-C) Frequencies of splenic cDC populations (panel B). Numbers (left) and frequencies (right) of splenic pDC populations (panel C). pDC = Lin^-^ CD11c^int^ B220^+^ Bst2^+^. Pooled from two independent experiments. (D) Representative flow plots (left) and frequencies (right) of migratory (CD11c^int^ MHC-II^hi^) and resident (CD11c^hi^ MHC-II^int^) cDC populations in skin-draining lymph nodes. Representative of at least three independent experiments. (E) Representative flow plots (left, in skin) and frequencies (right, in skin and skin-draining lymph nodes) of Langerhans cells (Lin^-^ CD11c^+^ MHC-II^+^ EpCAM^+^ CD207^+^ in skin, Lin^-^ CD11c^int^ MHC-II^hi^ EpCAM^+^ CD11b^+^ in skin-draining lymph nodes). Pooled from two independent experiments (skin). (F) Representative flow plots (left) and frequencies (right) of lung alveolar macrophages (CD64^+^ CD11c^hi^ Siglec-F^+^). Representative of two independent experiments. (G) Phenotypic characterization of splenic cDC2. Representative histograms (left) and MFI (right) of CD11b on Esam^+^ cDC2 from indicated mice. Pooled from two independent experiments. (H) Proportional histograms depicting overlap between Mat^ON^ and Notch signatures. (I) RNA-seq was performed on purified splenic cDCs, and GSEA was performed as in Figure 1E-F. Shown is GSEA of Notch signature (after removing overlapping Mat^ON^ signature) among differentially expressed genes (DEGs) between control and *Smc3*^Δ^ cDC1s. ES = enrichment score, NES = normalized enrichment score. Symbols represent individual mice, and bars represent mean. Control = *CD11c-*Cre (grey circles) and *Smc3^fl/fl^* (grey triangles), *Smc3*^ΔDC^ = *CD11c-*Cre *Smc3^fl/fl^* (blue circles) and *CD11c-*Cre *Smc3^fl/-^* (blue triangles). Statistical differences were evaluated using the Mann-Whitney test. **p < 0.01, ****p < 0.0001; n.s. not significant.

**Figure S2.**
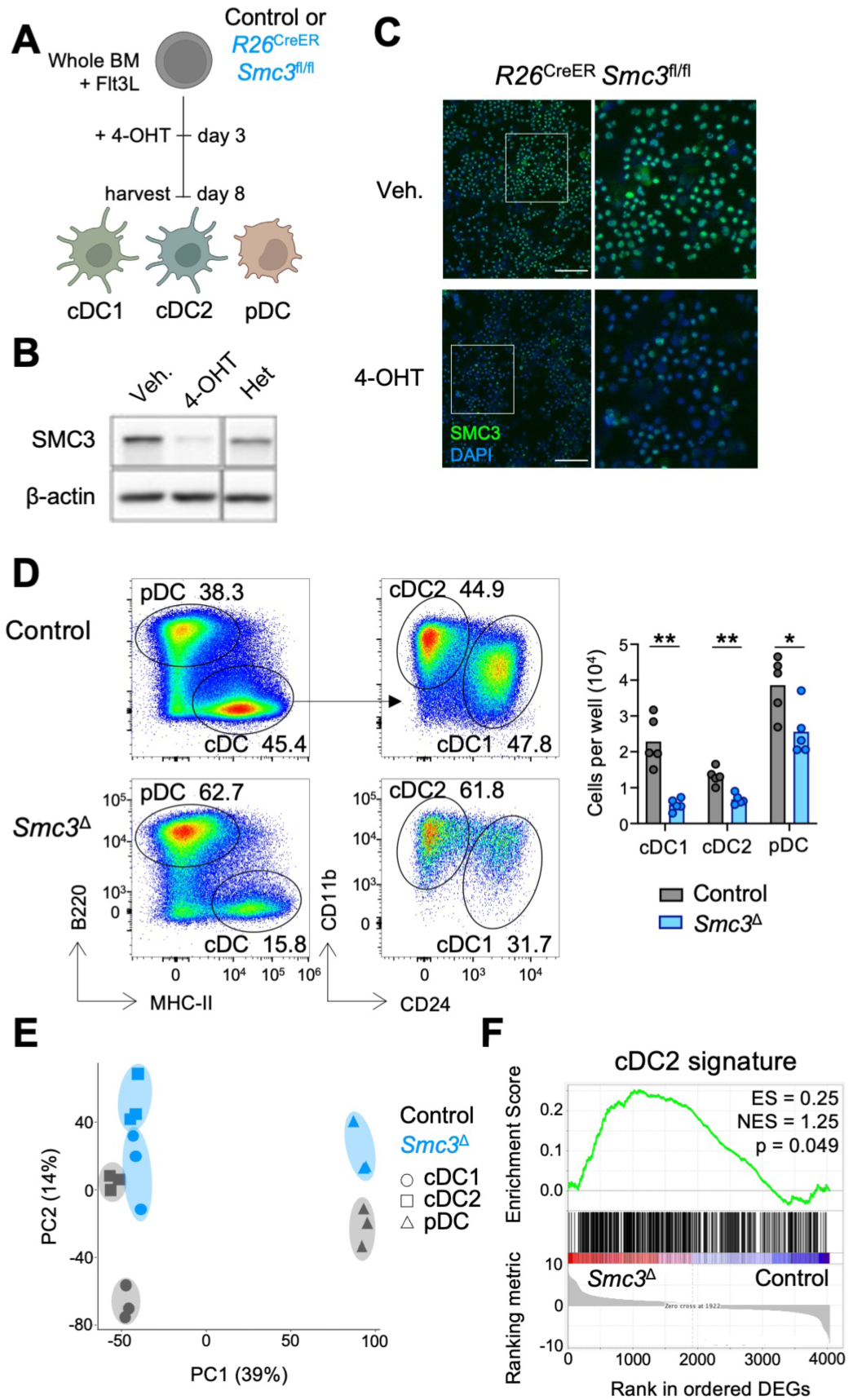
Cohesin is Required for *In Vitro* cDC Differentiation. (A) Schematic of experimental design. Bone marrow cells from *R26^CreER/^*^+^ (Control) or *R26^CreER/^*^+^ *Smc3^fl/fl^* (*Smc3*^Δ^) mice were differentiated into DCs (FL-BMDCs) *in vitro* in the presence of Flt3L. 4-hydroxytamoxifen (4-OHT) was added to cultures at day 3 to induce recombination of the floxed *Smc3* allele. DCs in the cultures were analyzed on day 8. (B) Western blot of SMC3 protein at differentiation day 8 in 4-OHT- or vehicle-treated *Smc3*^Δ^ cultures. *CD11c-*Cre *Smc3^fl/^*^-^ mice (Het) were used for comparison. Representative of two independent experiments. (C) Immunofluorescence staining of SMC3 protein. Right image is a zoom of inset on left image. Scale bar, 150 μm. (D) Representative flow plots (left) and numbers (right) of indicated DC subsets. cDC1 = CD11c^+^ MHC-II^+^ B220^-^ CD11b^int^ CD24^+^, cDC2 = CD11c^+^ MHC-II^+^ B220^-^ CD11b^hi^ CD24^-^, pDC = CD11c^+^ MHC-II^-^ B220^+^ Siglec-H^+^. (E-F) RNA-seq was performed on purified FL-BMDC subsets. PCA (panel E) and GSEA (panel F) of cDC2 signature (*37*) among differentially expressed genes (DEGs) between control and *Smc3*^Δ^ cDC1s. ES = enrichment score, NES = normalized enrichment score. Symbols represent individual mice, and bars represent mean. Statistical differences were evaluated using the Mann-Whitney test. *p < 0.05, **p < 0.01.

**Figure S3.**
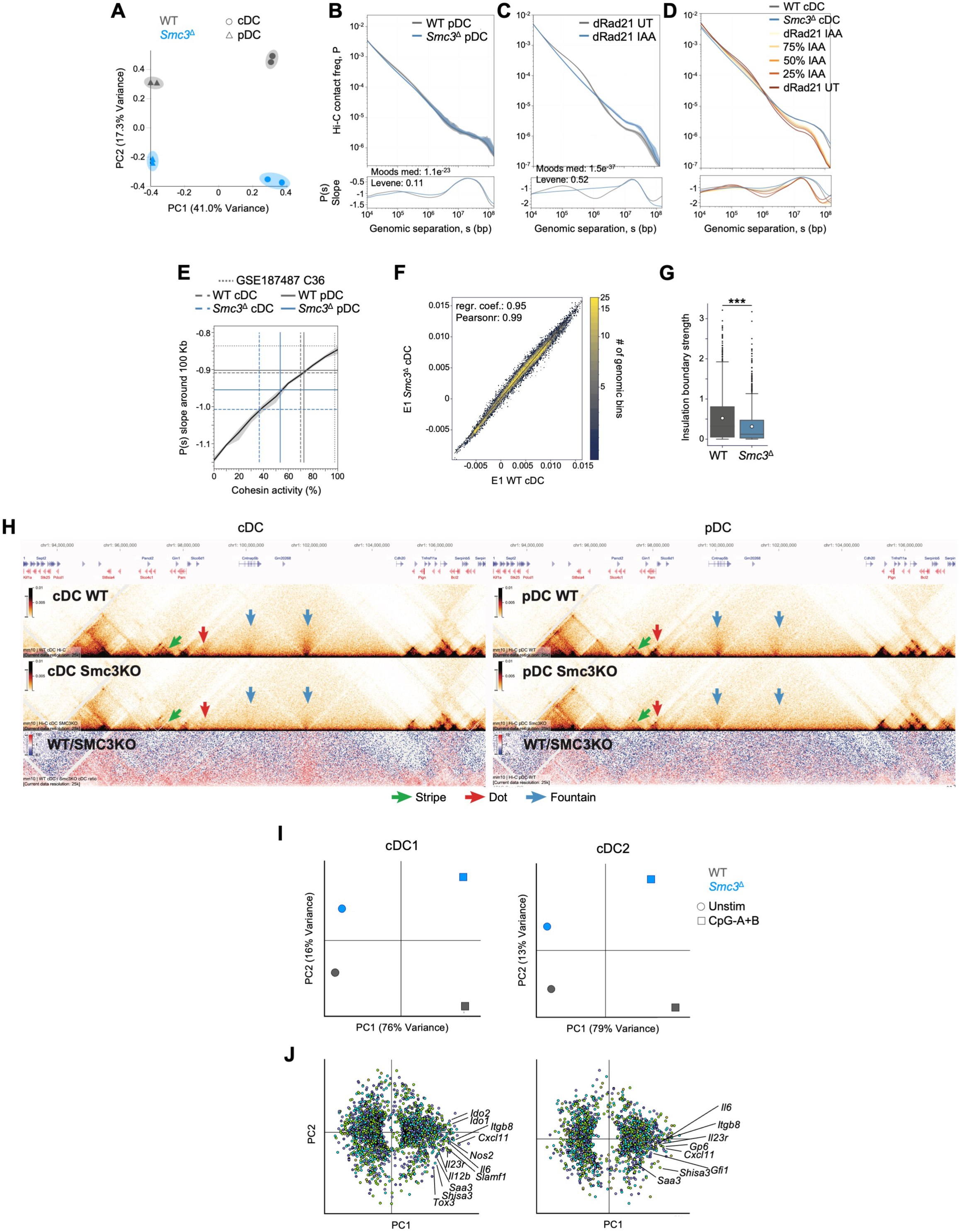
Cohesin Regulates Genome Organization in DCs. (A) PCA of Hi-C data, based on calculating pairwise stratum-adjusted correlation coefficients at 10 Kb resolution. (B) P(s) curves (top) and their slopes (bottom) for control and *Smc3*^Δ^ pDCs. Shaded regions represent the 90% confidence interval at each genomic separation range. Statistical differences between the distributions of slopes at 50-150 Kb were calculated with Moods median test and Levene test for variance equivalence. (C) P(s) curves (top) and their slopes (bottom) for auxin-treated (IAA) and untreated (UT) mESCs with degron-tagged Rad21 (*46*). (D-E) Estimate of cohesin activity in DC samples by comparing the P(s) slope drop at 100 Kb genomic separation in the samples with those of *in silico* mixtures of auxin-treated and untreated cells with degron-tagged Rad21 (*46*). (F) Scatterplot of E1 for each 25 Kb bin in control versus *Smc3*^Δ^ cDCs. (G) Insulation boundary strength in control versus *Smc3*^Δ^ cDCs. ***p < 0.001. (H) Contact frequency maps at a representative locus displaying a weakening of cohesin-dependent chromatin structures in *Smc3*^Δ^ relative to control cDCs (left) and pDCs (right). (I-J) cDC1s and cDC2s were purified for RNA-Seq from control and *Smc3*^Δ^ FL-BMDC cultures that were either unstimulated or stimulated for 6 hours with CpG-A and CpG-B. PCA analysis (panel I) and individual genes driving each PC (panel J) are shown. Genes are colored according to average expression (blue=low, orange=high).

**Figure S4.**
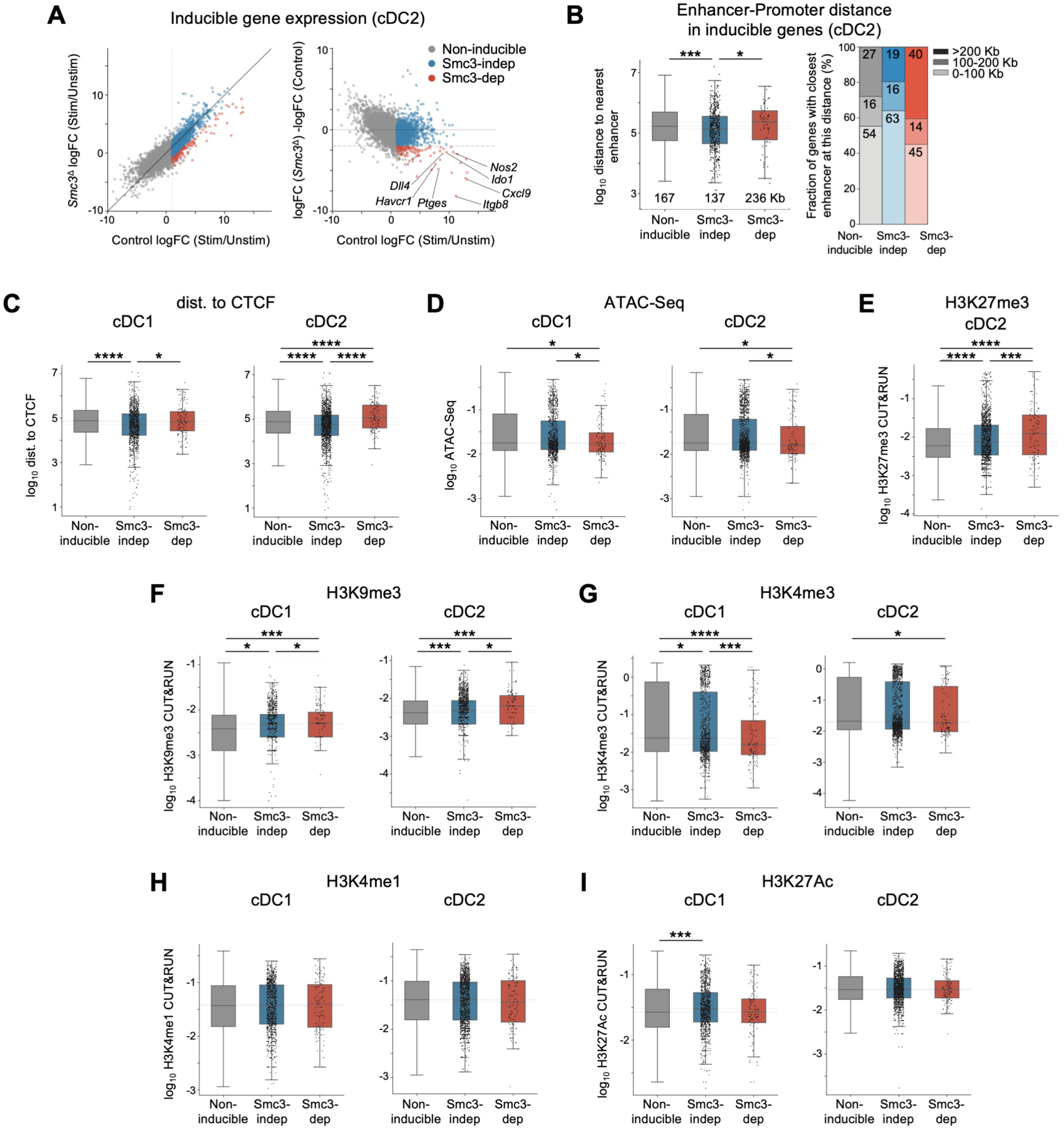
Chromatin Features of SMC3-Dependent Genes During cDC Activation. (A) Scatterplots of genes in control and *Smc3*^Δ^ cDC2s with or without CpG activation (as shown in Fig. 3F foe cDC1). Each gene is colored according to whether it is inducible and Smc3-dependent (red), inducible but Smc3-independent (blue) or non-inducible (grey). Select Smc3-dependent genes are indicated. (B) Distance of Smc3-dependent, Smc3-independent and non-inducible genes to their nearest enhancer in cDC2s (left). Dashed lines represent the medians of the distributions, and the median value in Kb is indicated on the plot. Statistical differences were calculated with the Mann-Whitney one-sided test. Proportion of indicated gene subsets that have their nearest enhancer located within 100 Kb, 100-200 Kb or more than 200 Kb away from their transcription start site (right). (C-I) Distance to the nearest ChIP-seq-derived CTCF binding site (C); magnitude of ATAC-seq derived chromatin accessibility signal (D); and CUT&RUN-derived H3K27me3 (E), H3K9me3 (F), H3K4me3 (G), H3K4me1 (H), and H3K27Ac (I) signal in cDCs within the insulated neighborhood of Smc3-dependent, Smc3-independent and non-inducible genes during cDC1 or cDC2 activation. Dashed lines represent the medians of the distributions. Statistical differences were calculated with the Mann-Whitney one-sided test. Data are presented as box plots. *p < 0.05, ***p < 0.001, ****p < 0.0001.

**Figure S5.**
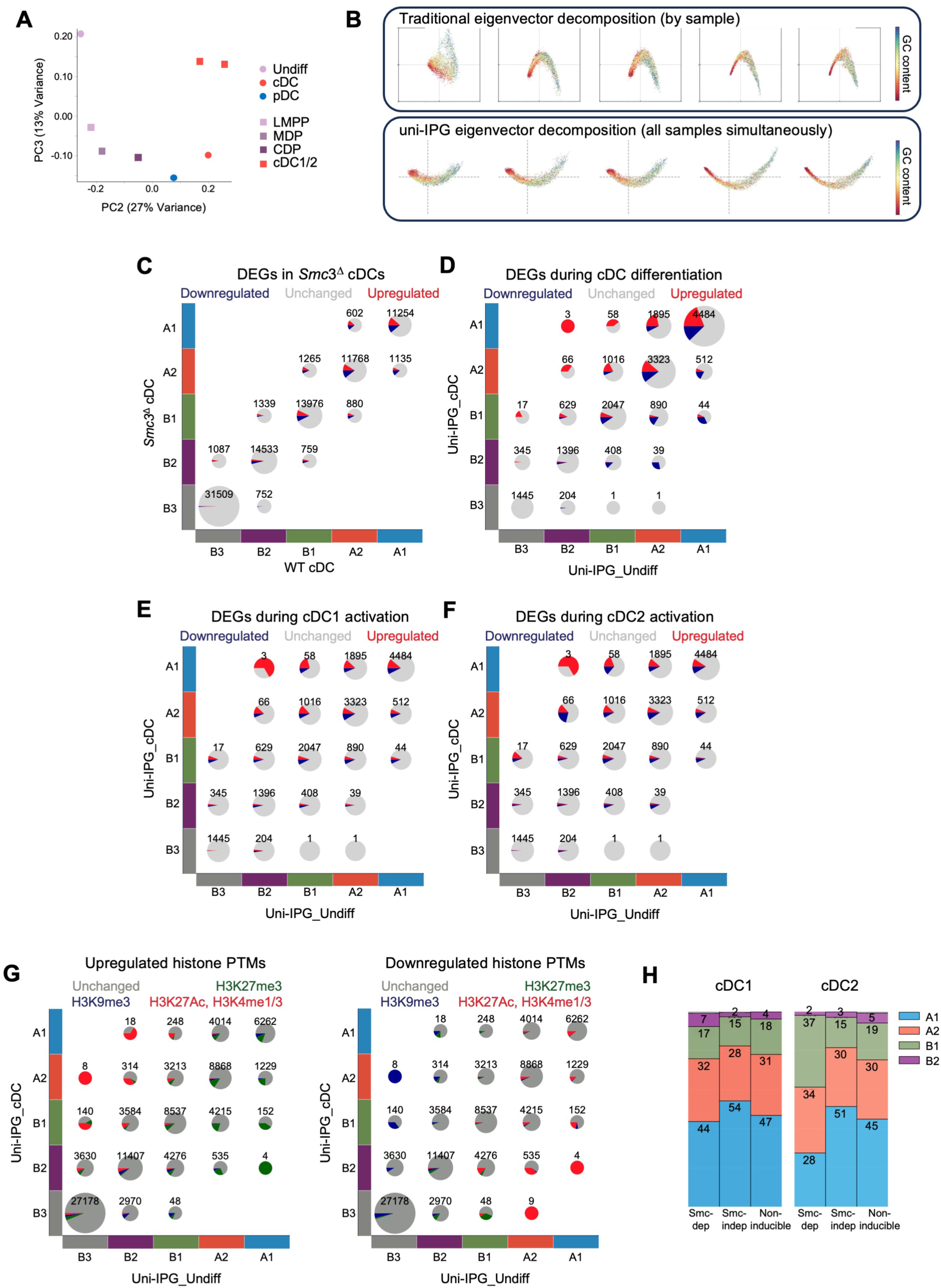
Additional Characterization of Genome Organization Remodeling During DC Differentiation. (A) PCA of the Hi-C data from HoxB8-FL cell differentiation (Fig. 4A) along with the data from *ex vivo* DC progenitors (*54*), based on calculating stratum-adjusted correlation coefficients on condition merged Hi-C maps at 10 Kb resolution. LMPP = lymphoid-primed multipotent progenitors, MDP = monocyte-DC progenitor, CDP = common DC progenitor. (B) Comparison of eigenvector decomposition using traditional “by sample” (top) or “simultaneous” uni-IPG approach (bottom). Cell types (left to right): progenitors, cDCs and pDCs from HoxB8-FL cells, cDCs and pDCs from FL-BMDC cultures. (C) Dot plot of IPG states in control and *Smc3*^Δ^ cDCs. Highlighted in red and blue are genes upregulated or downregulated respectively in *Smc3*^Δ^ cDCs. Number of bins in each pair of states is indicated. (D-F) Dot plots of IPG transitions between progenitors and cDCs. Highlighted in red and blue are genes upregulated or downregulated respectively in cDCs during differentiation (panel D), cDC1s after CpG stimulation (panel E), and cDC2s after CpG stimulation (panel F). Number of genes in each pair of states is indicated. (G) Dot plots of IPG transitions between progenitors and cDCs. Highlighted are histone modifications upregulated (left) or downregulated (right) in each pair of IPG transitions. Number of bins in each pair of states is indicated. (H) Uni-IPG composition of Smc3-dependent, Smc3-independent and non-inducible genes in cDC1s (left) and cDC2 (right) prior to CpG stimulation.

**Figure S6.**
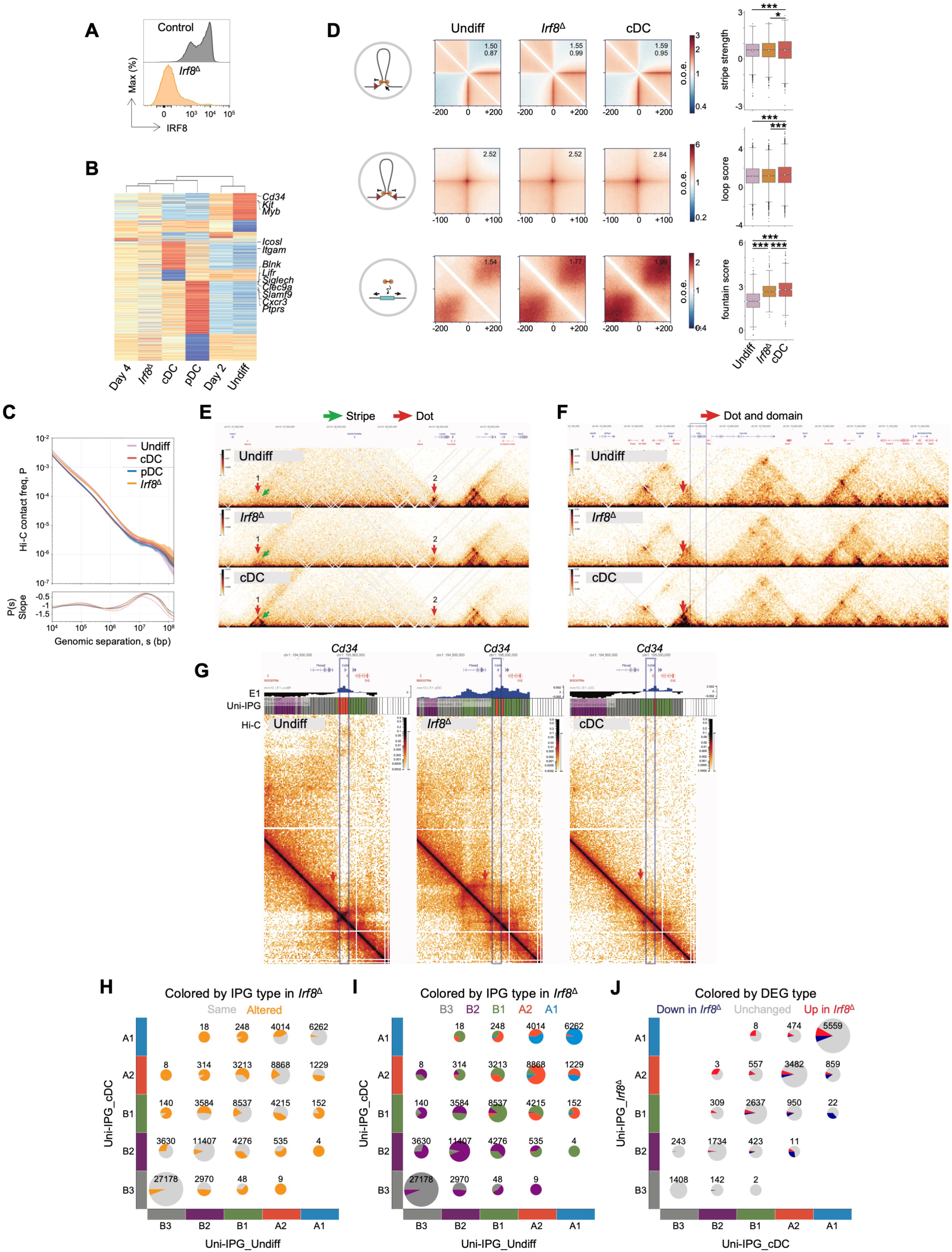
Additional Characterization of IRF8-Mediated Chromatin Effects in Differentiating DCs. An sgRNA targeting *Irf8* was used to generate IRF8-deficient (*Irf8*^Δ^) HoxB8-FL cells using Cas9-RNPs. (A) Histograms of IRF8 staining in CD11c^+^ cells differentiated from *Irf8*^Δ^ progenitors or progenitors electroporated with a non-targeting control sgRNA. Representative of two independent experiments. (B) RNA-seq was performed on HoxB8-FL cells at several stages along their differentiation trajectory from undifferentiated progenitors (Undiff) to mature DCs as well as on *Irf8*^Δ^ DCs. Samples are hierarchically clustered based on transcriptional similarities. Marker genes characteristic for each developmental timepoint are indicated on the heatmap. (C) P(s) curves (top) and their slopes (bottom) for progenitors, cDCs, pDCs and *Irf8*^Δ^ DCs. Shaded regions represent the 90% confidence interval at each genomic separation range. (D) Average pileups around forward-oriented CTCF peaks (stripes, top row), around pairs of CTCF peaks with convergent orientation (dots, middle row) and at active chromatin (fountains, bottom row) (left) and summary statistics of their strength (right) in progenitors, cDCs and *Irf8*^Δ^ DCs. Summary statistics are presented as box plots. Statistical differences were calculated with the Mann-Whitney one-sided test. *p < 0.05, ***p < 0.001. (E-F) On-diagonal Hi-C maps in HoxB8-FL progenitors as well as cDCs and *Irf8*^Δ^ DCs. (G) Hi-C maps in the vicinity of the *Cd34* locus in HoxB8-FL progenitors as well as cDCs and *Irf8*^Δ^ DCs. First projected spectral component (E1) and uni-IPG tracks are shown above the Hi-C maps. (H-I) Dot plots of IPG transitions between progenitors and cDCs. Highlighted in orange are altered IPGs in *Irf8*^Δ^ DCs (panel H) and the actual IPG type in *Irf8*^Δ^ DCs (panel I). Number of bins in each pair of states is indicated. (J) Dot plot of IPG state in cDCs and *Irf8*^Δ^ DCs. Highlighted in red and blue are genes upregulated and downregulated respectively in *Irf8*^Δ^ DCs relative to cDCs. Number of genes in each pair of states is indicated.

**Figure S7.**
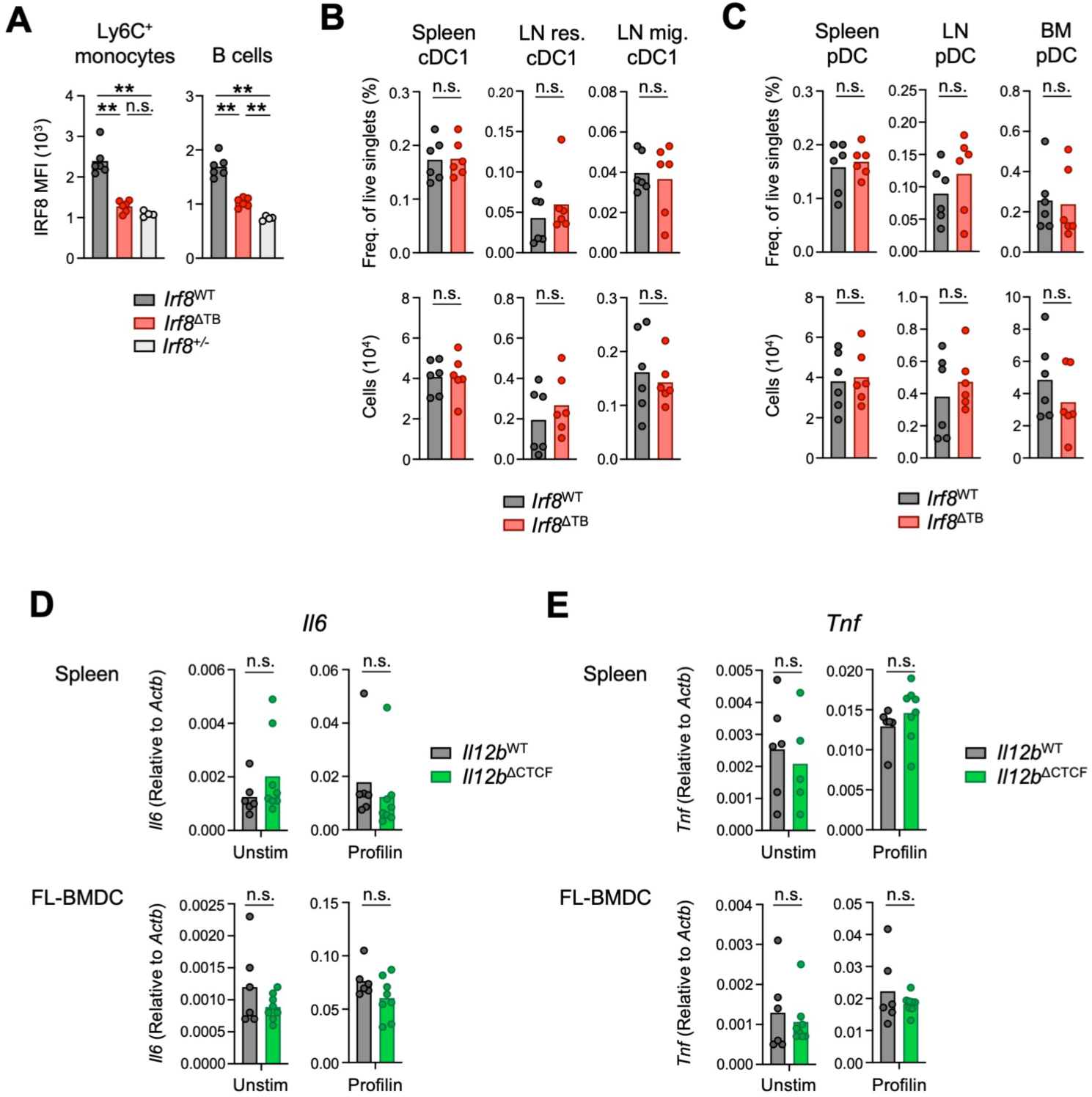
Additional characterization of *Irf8*^ΔTB^ and *Il12b*^ΔCTCF^ mice. (A-C) Additional characterization of *Irf8*^ΔΤΒ^ mice (A) Quantification of IRF8 MFI in splenic Ly6C^+^ monocytes and B cells. *Irf8^+/-^* mice were used as controls with 50% reduction of IRF8 expression. (B-C) Frequencies (top) and numbers (bottom) of cDC1 (panel B) and pDC (panel C) in indicated tissues. (D-E) Additional characterization of *Il12b*^ΔCTCF^ mice. Shown is RT-qPCR for *Il6* (panel D) and *Tnf* (panel E) (expressed as a ratio relative to that of the housekeeping gene *Actb*) in spleen enriched for cDC1 (top) and cDC1s purified from FL-BMDC cultures (bottom) of indicated mice that were either unstimulated (left) or stimulated with profilin (right) for 4 hours. Symbols represent individual mice, and bars represent mean. Statistical differences were evaluated using the Mann-Whitney test. **p < 0.01; n.s. not significant.

**Supplemental Methods Figure 1.**
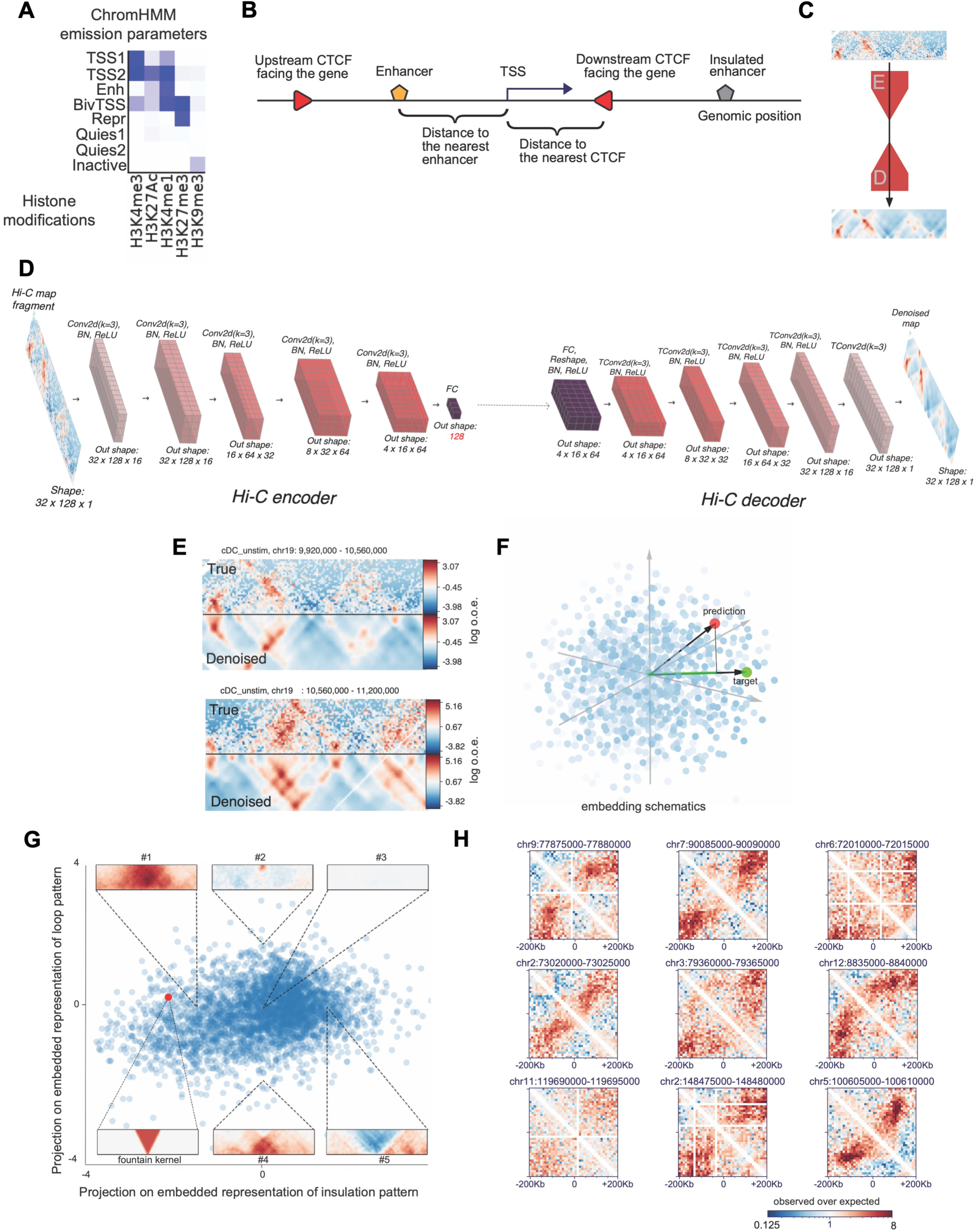
Methods for the analysis of chromatin states. (A) ChromHMM emission parameters for training on the available CUT&RUN data. Shown is the output of ChromHMM tool representing a heatmap of histone modification intensities across the chromatin states as described in the Methods: TSS1/2, active transcription states; Enh, a state with enriched H3K4me1; BivTSS, a bivalent transcription state enriched in both H3K27me3 and H3K4me1;Repr, a repressed state enriched in H3K27me3; Quies1/2, quiescent chromatin with all histone marks depleted; Inactive, a state enriched in H3K9me3. (B) Schematic of the definition of the closest enhancer to the gene in the insulated neighborhood. (C-H) Autoencoder-based fountain calling. (C) Schematic of the process: Hi-C matrices are encoded for noise reduction and embedding different genomic regions in the same space. E, encoder; D, decoder representing the neural network blocks. The arrow indicates the information flow through the neural network. (D) Schematic of the neural network architecture for the autoencoder-based encoding of Hi-C matrices from panel C. “Conv2d” are 2d-convolutional blocks, “FC” are fully connected blocks, “TConv2d” are transposed convolutional blocks, “BN” are batch-normalization layers, and ReLU is rectified linear unit. (E) Two randomly selected examples of loci before and after de-noising with the autoencoder. (F) Schematic of the calculation of fountain score. The red dot is an embedded representation of the position in the embedded space for the genomic region, and the green dot is an embedded representation of the target structure — the fountain. For a given region, the projection is calculated between the vector of embedded representation (prediction) and the fountain (target) vector. The value of this projection is interpreted as a fountain score for detecting reliable peaks in fountain score tracks and, eventually, fountains (see Methods). (G) A representation of embedded space of the autoencoder as a projection to representations of loop and insulation patterns. Blue dots represent individual genomic regions, with typical decoded maps illustrated for several regions. Red dot represents a region encoding a fountain. (H) Representative examples of fountains identified by the autoencoder.

